# Lysosome docking to Omegasomes and ER-connected Phagophores Occurs During DNAJB12 and GABARAP-Dependent Selective Autophagy of Misfolded P23H-Rhodopsin

**DOI:** 10.1101/2021.10.20.465144

**Authors:** Andrew Kennedy, Hong Yu Ren, Victoria J. Madden, Douglas M. Cyr

## Abstract

We report on how the ER-associated-autophagy pathway (ERAA) delivers P23H-rhodopsin (P23H-R) to the lysosome. P23H-R accumulates in an ERAD-resistant conformation that is stabilized by DNAJB12 and Hsp70. P23H-R, DNAJB12, and FIP200 co-localize in discrete foci that punctuate the rim of omegasome rings coated by WIPI1. P23H-R tubules thread through the wall of WIPI1 rings into their central cavity. Transfer of P23H-R from ER-connected phagophores to lysosomes requires GABARAP, and is associated with the transient docking of lysosomes to WIPI1 rings. Instances of lysosomes docking to WIPI1 foci are constitutive, and increase 250% upon P23H-R expression. After departure from WIPI1 rings, new patches of P23H-R are seen in the membranes of lysosomes. The absence of GABARAP prevents transfer of P23H-R from phagophores to lysosomes without interfering with docking. These data identify lysosome docking to omegasomes as an important step in the DNAJB12 and GABARAP-dependent autophagic disposal of dominantly toxic P23H-R.

## Introduction

The endoplasmic reticulum (ER) is the site where biogenesis of membrane proteins initiates in a process that requires the coordinated folding and assembly of subdomains that are located in the cytosol, lumen, and ER-membrane (Phillips and Miller, 2021). This process can be inefficient for both wild-type (WT) and disease proteins, which challenge ER protein quality control (ERQC) systems with the accumulation potentially toxic misfolded intermediates (Hipp et al., 2019). For example, retinitis pigmentosa arises from missense mutations in the light sensor rhodopsin that lead to its misfolding and retention in the ER. Rhodopsin is a G-protein coupled receptor that contains a conserved disulfide whose formation is required for binding of the light sensing cofactor 9-cis-retina (Athanasiou et al., 2018). Folding of rhodopsin, therefore, involves the initial collapse of its glycosylated nascent chains, disulfide bond formation, and co-factor binding. This stepwise process is facilitated by the glycan binding and ER-transmembrane molecular chaperone calnexin, as well as cytosolic Hsp40 and Hsp70 chaperone pairs that function on the cytoplasmic face of the ER (Chapple and Cheetham, 2003; Saliba et al., 2002).

Intrinsic inefficiencies in rhodopsin assembly lead kinetically-trapped intermediates to accumulate in the ER-membrane where they are selected for premature degradation by the ER-associated proteasomal degradation pathway, ERAD(Chiang et al., 2012). The majority of nascent misfolded membrane proteins are cleared via ERAD involving their ER-retention, ubiquitination, retrograde translocation to the cytosol, and proteasomal degradation(Ferro-Novick et al., 2021). Misfolded rhodopsin is a client of the ERAD pathway, but some rhodopsin mutants causing retinitis pigmentosa, such as P23H-rhodopsin (P23H-R), are inefficiently degraded by ERAD (Meng et al., 2020). Resultant accumulation of dominantly toxic conformers of P23H-R causes retinal degeneration and blindness (Athanasiou et al., 2018). Therapeutic suppression of mutant rhodopsin proteotoxicity remains to be discovered, and there are no drugs available to treat the underlying cause of retinitis pigmentosa(Meng et al., 2020).

To develop approaches that alleviate protein misfolding diseases such a retinitis pigmentosa, the identification of cellular mechanisms that buffer the accumulation of toxic protein species in the ER-membrane system and secretory pathway is essential. Recent studies have identified selective-autophagic mechanisms that operate in unstarved cells that clear misfolded and ERAD-resistant protein intermediates from the ER-lumen, ER-exit sites, and ER-membrane (Ferro-Novick et al., 2021; Wilkinson, 2019a, b). ERAD-resistant forms of misfolded α-1-antitrypsin mutants are cleared from the ER-lumen by complexes containing calnexin and the ER-autophagy (ER-phagy) receptor FAM134B (Fregno and Molinari, 2019). Misfolded proteins that enter ER-exit sites but cannot be secreted, are degraded by micro-ER-phagy in a FAM134b-dependent process. This is thought to involve vesicle-mediated movement of misfolded and ERAD-resistant proteins to lysosomes for degradation (Loi and Molinari, 2020). ERAD-resistant misfolded membrane proteins accumulating in ER-membranes are degraded via a pathway that is independent of FAM134B but requiring the transmembrane Hsp40 DNAJB12 (JB12) and cytosolic Hsp70 (He et al., 2021; Houck et al., 2014).

Studies on the Gonadotropin Hormone Releasing Hormone Receptor (GnRHR), a G-protein coupled receptor (GPCR), demonstrated that the mutation E90K causes folding defects that occur downstream of the formation of the conserved disulfide which mediates formation of its ligand binding site (Houck et al., 2014). This causes ERAD-resistant pools of E90K-GnRHR to accumulate in the ER and stimulates flux through the JB12/Hsp70-dependent pathway for selective autophagy (Houck et al., 2014). Kinetically-trapped and ERAD-resistant intermediates of misfolded CFTR have also been shown to be cleared from the ER via the selective autophagy process requiring JB12 and Hsp70 (He et al., 2021). Kinetically-trapped intermediates of membrane proteins that are resistant to ERAD (ERAD-RMPs) accumulate in ER-membranes because they cannot be unfolded and retrotranslocated into to the cytosol. This invites them to remain associated with JB12 and Hsp70 for extended times, leading to downstream association of JB12 and Hsp70 with autophagy initiation machinery. These events are proposed to drive the metamorphosis of ER tubules containing ERAD-RMPs into phagophores (autophagosome precursors) within omegasomes (autophagy initiation sites) that are decorated with the phosphatidylinositol 3-phosphate (PtdIns3*P*) binding protein WIPI1 (WD-repeat protein interacting with phosphoinositide’s)(He et al., 2021). Yet, it is presently not clear how ERAD-RMPS are packaged into autophagosomes and transferred from the ER to lysosomes(Ferro-Novick et al., 2021).

Herein, we report on the study of the mechanisms for the triage by the JB12/Hsp70 chaperone complex of WT rhodopsin (WT-R) and P23H-R for degradation via ERAD or selective-autophagy within the ER-membrane system. This process does not require ER-phagy receptors (He et al., 2021), and is an alternative to the ERAD pathway, so it is referred to as ER-associated autophagy (ERAA). Both WT-R and P23H-R were detected in complexes with JB12 and Hsp70, with levels of P23H-R:JB12 complexes being dramatically higher than those with WT-R. P23H-R accumulates in ER tubules with JB12 that interact with components of omegasome rings DFCP1 (double FYVE domain-containing protein) and WIPI1 that bind PtdIns3*P* produced by the Beclin-1:VPS34 autophagy initiation kinase(Ferro-Novick et al., 2021). P23H-R, but not WT-R, also accumulated in lysosomes with ATG8 family members and the lysosome marker LAMP. Interference with the folding of P23H-R via mutation of the highly conserved cysteines that form rhodopsin’s disulfide prevented P23H-R entry into omegasomes, so the conformation of misfolded rhodopsin is fate-determining. ER tubules enriched in P23H-R containing GABARAP extend into the cytosol from WIPI1 decorated rings of ER-membrane. Lysosomes transiently dock with WIPI1 rings and then become enveloped by P23H-R containing phagophores. Lysosomes are detected in association with 10% of all cellular WIPI1 rings. P23H-R expression stimulates the association of lysosomes with WIPI1 rings 2.5-fold. Lysosome docking to WIPI1 rings occurs for 10-60 seconds and after departure patches of P23H-R are detected in the membranes of lysosomes. The depletion of GABARAP prevents transfer of P23H-R from the ER to lysosomes, but does not hinder the sequestration of P23H-R into phagophores or lysosome docking to WIPI1 rings. Models for the sequential roles of JB12 and Hsp70, GABARAP, and lysosome docking to WIPI1 rings in the clearance of ERAD-resistant and dominantly toxic pools of P23H-R from the ER-tubular network by ERAA will be discussed.

## Results

### Triage of P23H-Rhodopsin Between Proteasomal and Lysosomal Degradation is Conformation Dependent

To explore the mechanism for triage in the ER of misfolded rhodopsin intermediates, we evaluated the fate of WT-R and P23H-R through microscopy and biochemical methods (Figure 1). WT-R and P23H-R behave similarly to WT-R-eGFP and P23H-R-eGFP in biochemical and cell biological assays that monitor their biogenesis (Chapple and Cheetham, 2003; Saliba et al., 2002). We therefore employed eGFP-tagged forms of WT-R and P23H-R in our studies and from here on refer to the eGFP-tagged proteins as WT-R and P23H-R.

**Figure 1.**
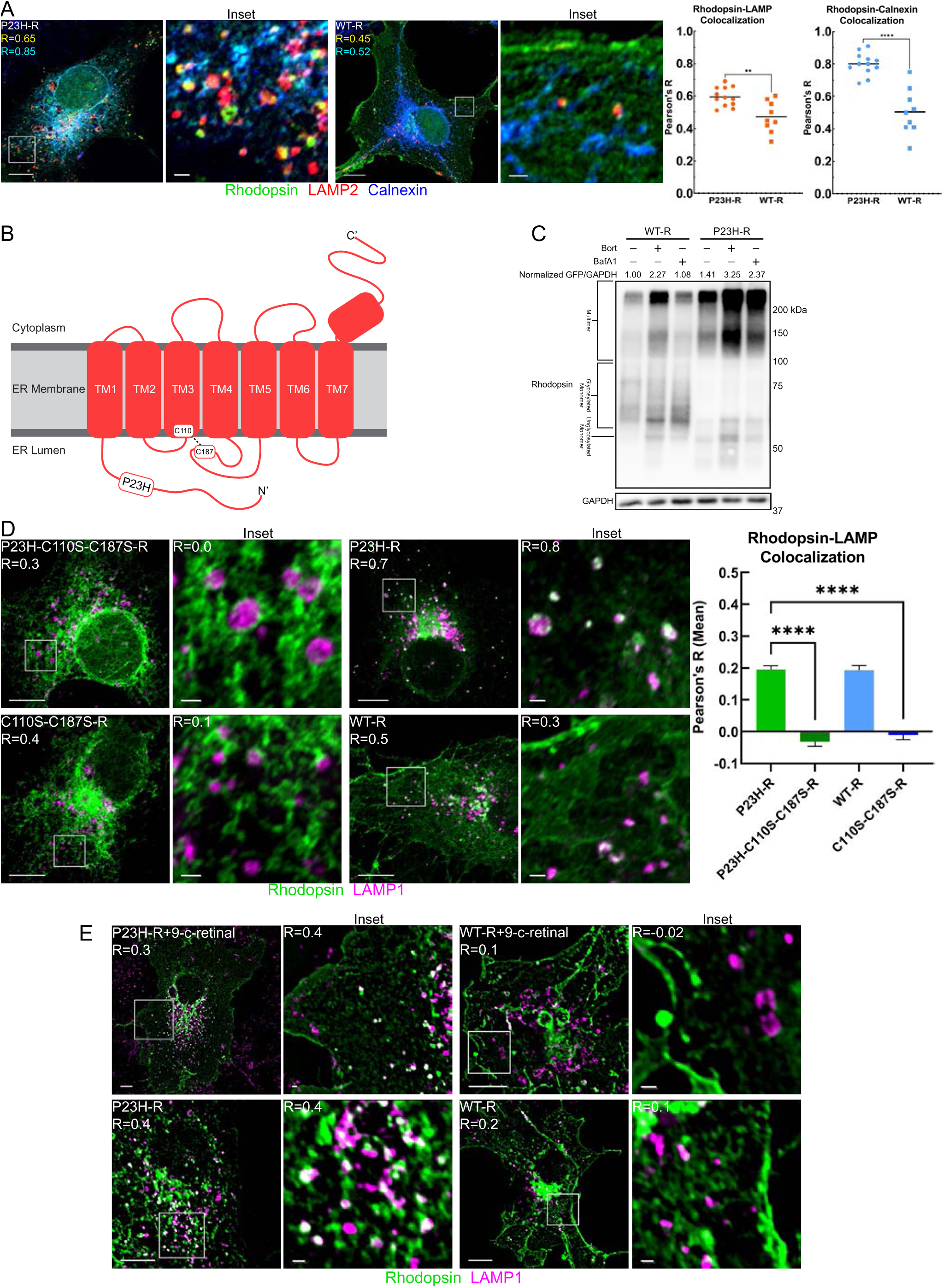
P23H-Rhodopsin Triage Between the Proteasome and Lysosome. COS-7 cells transiently transfected with eGFP-Rhodopsin for WT or indicated mutations. All microscopy conducted in fixed cells with immunofluorescence staining for indicated marker. (A) Left: Fixed-cell confocal microscopy on Zeiss 880 with Airyscan - LAMP2a (Red), Rhodopsin (Green), and Calnexin (Blue). Pearson’s R values indicated in whole-cell panels by color - top number (yellow) LAMP-Rho; bottom number (cyan) Calnexin-Rho Right: Dot plots of Pearson’s R with unpaired t-test - line indicates mean (B) Secondary structure of Rhodopsin (C) Western blot of COS-7 cells expressing either WT-R or P23H-R and treated with either 10µM Bortezomib (Bort) or 100nM Bafilomycin-A1 (BafA1) for 6 hours. (D) Left: Fixed-cell confocal microscopy of indicated Rhodopsin mutant on Zeiss 880 with Airyscan – Rhodopsin (Green), LAMP1 (Magenta). R indicates Pearson’s R for Rhodopsin-LAMP1 colocalization Right: Bar graph of mean Pearson’s R values between Rhodopsin and LAMP1 calculated within identified LAMP1 objects. Kruskal-Wallis and Dunn’s multiple comparisons tests (n=1240). Error bars for SEM (E) Fixed-cell epifluorescent microscopy of WT-R or P23H-R ± 10µM 9-cis-retinal–Rhodopsin (Green), LAMP1 (Magenta). R indicates Pearson’s R for Rhodopsin-LAMP1 colocalization \*\*\*\**P* < 0.0001 Scale bars: whole cell views = 10µm; insets = 1µm

To investigate mechanisms for triage of rhodopsin intermediates we used experimental conditions where WT-R and P23H-R were expressed from 50ng of the same expression plasmid used in previous studies that suggested P23H-R is aggregation prone and becomes sequestered within aggresomes (Saliba et al., 2002). Under these conditions WT-R and P23H-R are biosynthesized at low levels that are within the cellular capacity for their biogenesis. WT-R accumulates at the plasma membrane (PM) and in low levels with the ER-membrane with calnexin. In contrast, P23H-R co-localizes in primarily with calnexin and the lysosome associated membrane protein 2 (LAMP2) (Figure 1A) (LAMP1 and LAMP2 are both lysosome makers and recognized by different mouse monoclonal anti-bodies, so they are used interchangeably, and from here on are denoted as LAMP). Since WT-R is endocytosed from the PM, a pool of it is detected in lysosomes, but co-localization of P23H-R with LAMP occurs at a higher degree than for WT-R (Figure 1A). Since P23H-R is retained in the ER, its route for traffic to the lysosome is also different from the route for WT-R.

To biochemically evaluate the route for degradation of WT-R and P23H-R we tested the sensitivity of their steady-state accumulation to the proteasome inhibitor bortezomib (Bort) and the lysosomal proton pump inhibitor bafilomycin A1 (BafA1). BafA1 causes increased pH of the lysosome lumen and thereby renders lysosomal endoproteases inactive. WT-R and P23H-R run on SDS-PAGE gels with different mobilities that correspond to a faster glycosylated ER pool, a slower heavily glycosylated form trafficked to the PM, or multimers that form after cell lysis. Since WT-R and P23H-R self-associate and migrate as higher molecular weight oligomers during SDS-PAGE, we quantitated the signals for the entire lane on western blots of cell lysates. Folding of WT-R is inefficient with misfolded intermediates being degraded by ERAD (Chapple and Cheetham, 2003; Saliba et al., 2002). Incubation of cells with Bort for 6-hours prior to harvest caused a greater than 2-fold increase in WT-R accumulation, but BafA1 did not increase WT-R accumulation. Bort acted to increase P23H-R accumulation around 2-fold as did BafA1 (Figure 1C). These data are consistent with imaging data showing P23H-R accumulation in both the ER with calnexin and in lysosomes with LAMP. These results are also consistent with previous studies on the triage of misfolded membrane proteins showing that the conformation of membrane protein intermediates is scanned on an individual basis for triage to pathways for proteasomal or lysosomal degradation (Buchberger, 2014; Chiang et al., 2012; He et al., 2021; Houck et al., 2014).

Models for membrane protein triage suggest that pools of ER-arrested intermediates contain globally misfolded conformers that are retrotranslocated for ERAD. There is also a population of kinetically trapped intermediates containing stable tertiary structure causing resistance to retrotranslocation and triage for lysosomal degradation (He et al., 2021; Houck et al., 2014). The cysteine residues C110 and C187 in rhodopsin form a highly conserved and stabilizing disulfide in GPCR biogenic intermediates (Conn et al., 2007). We therefore asked if destabilization via mutating C110 and C187 to serine would impact the degradation fate of P23H-R. C110S-C187S-R did not traffic to the cell surface, indicative of it having a non-native conformation. P23H-C110S-C187S-R showed a reduced propensity to accumulate in lysosomes with LAMP when compared to P23H-R (Figure 1D). To verify this observation, we employed object-based identification of LAMP vesicles (n=1240) with subsequent correlation coefficient calculations (Pearson’s R) between LAMP1 and the indicated rhodopsin mutant. Mean Pearson’s R values were significantly different when comparing P23H-C110S-C187S-R (mean R = −0.004) and P23H-R (mean R = 0.36), confirming a dramatic reduction in traffic to lysosomes. This is the case because P23H-C110S-C187S-R is globally misfolded and only accumulates in the ER-membrane system.

To evaluate how the stabilization of P23H-R impacts triage decisions, 9-cis-retinal (9CR), the chromophore of rhodopsin known to act as a pharmacologic chaperone (Meng et al., 2020), was included for 6-hours in cell culture media prior to fixation. Imaging revealed that the presence of 9CR in cell cultures promotes cell-surface localization of a fraction of P23H-R, indicating suppression of misfolding. P23H-R and LAMP1 colocalization was still detected in 9CR treated cells (Figure 1E). 9CR binding appears to stabilize ER pools of P23H-R, but cannot prevent ERAA-targeted conformers from forming. These collective data suggest that kinetically trapped intermediates of P23H-R which have stable, but non-native structure, are triaged for lysosomal degradation.

### Lysosomal Degradation of P23H-Rhodopsin is Facilitated by the ER-Associated Chaperones JB12 and Hsp70

JB12 and cytosolic Hsp70 dynamically interact with different ERQC machineries during the conformation-dependent triage of membrane proteins between assembly into active conformers, or degradation via the proteasome/lysosome (Grove et al., 2011; He et al., 2021; Houck et al., 2014; Li et al., 2017; Walczak et al., 2014). To determine the requirements for JB12 in the conformation-dependent triage of rhodopsin intermediates, the localization of endogenous JB12 with P23H-R or P23H-C110S-C187S-R was imaged. Both P23H-R and the destabilized P23H-C110S-C187S-R co-localized with JB12 in the ER (Figure 2A). Notably, the P23H-R signal that co-localized with JB12 was also seen in connected ER-membranes containing P23H-R, but devoid of JB12. This arrangement was not observed with P23H-C110S-C187S-R. These data suggest that P23H-R is segregated within the ER into JB12-free and JB12-bound pools, and below we define the importance of this segregation event in triage of P23H-R.

**Figure 2.**
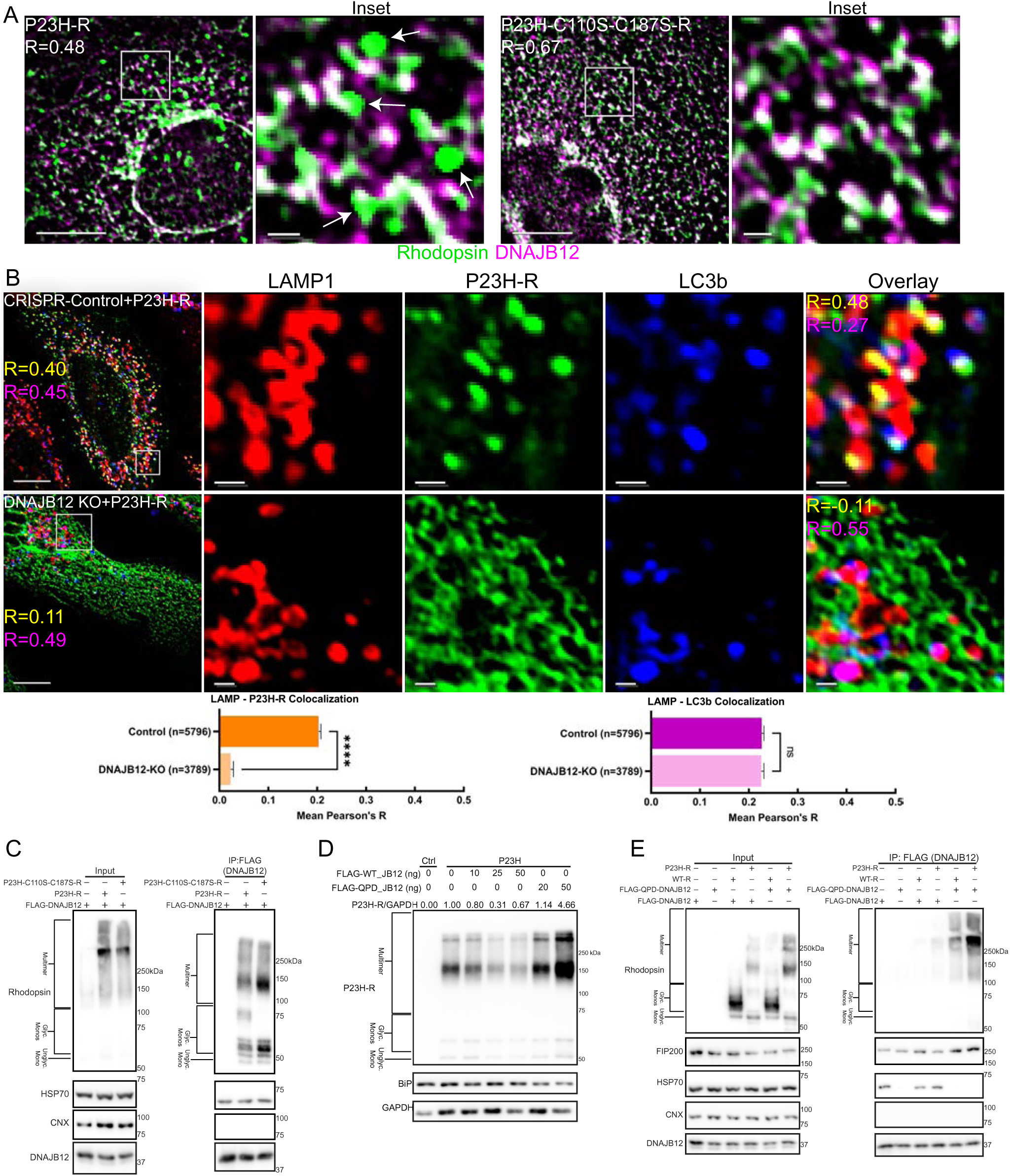
DNAJB12 and HSP70 Participate in the Triage of Misfolded P23H-Rhodopsin Towards Lysosomal Degradation. (A) Fixed-cell epifluorescent microscopy of COS-7 cells transiently transfected with P23H-R or P23H-C110S-C187S-R. Green-Rhodopsin; Magenta – DNAJB12, white arrows – P23H-R foci extending from the ER tubules containing DNAJB12. R indicates Pearson’s R for Rhodopsin-DNAJB12 colocalization (B) TOP: Fixed-cell epifluorescent microscopy of HeLa cell lines transduced with either a control (CRISPR-Control) or DNAJB12 knockout (CRISPR DNAJB12 KO) lentivirus, then transiently transfected with P23H-R. Immunofluorescence conducted for LAMP1 (Red) or LC3b (Blue). R values: top (yellow) P23H-R/LAMP1; bottom (Magenta) LAMP1/LC3b. BOTTOM: Bar graph of mean Pearson’s R values between Rhodopsin and LAMP1 (left) or LAMP1 and LC3b (right) calculated within identified LAMP1 objects. Mann-Whitney test - error bars for SEM (C) Western blot of FLAG pull-down in COS-7 cells transiently transfected with FLAG-DNAJB12 and either P23H-R or P23H-C110S-C187S-R (D) Western blot of COS-7 cells transiently transfected with P23H-R and increasing concentrations of either FLAG-DNAJB12 (WT) or FLAG-QPD-DNAJB12 (DNAJB12 mutant with dominant-negative HSP70 interaction domain) (E) Western blot of FLAG pull-down in HeLa cells transiently transfected with P23H-R or WT-R and either FLAG-DNAJB12 or FLAG-QPD-DNAJB12 \*\*\*\**P* < 0.0001 Scale bars: whole cell views = 10µm; insets = 1µm

To determine if JB12 is an integral participant in the P23H-R triage mechanism, a knockout cell line (JB12-KO) was generated using the CRISPR-Cas9 system in HeLa cells. Depletion of JB12 was confirmed by western blot (Figure S2A) and immunostaining in fixed-cell microscopy (Figure 2B). CRISPR-Control and JB12-KO knockdown cells were transiently transfected with P23H-R and imaged. JB12 knockout impaired accumulation of P23H-R in lysosomes (Figure 2B). Object-based image analysis of LAMP vesicles in CRISPR-control cells (n=5796) and JB12-KO (n=3789) reveals that the loss of JB12 reduced P23H-R presence in lysosomes (Figure 2B). Importantly, endogenous LC3b retained its localization to the lysosome, indicating the loss of DNAJB12 did not hinder basal autophagic flux(Figure 2B). The loss of JB12 does not hinder basal autophagic flux, but specifically reduces accumulation of P23H-R in lysosomes.

It remained unclear if JB12 influenced P23H-R triage directly or indirectly, so their presence in complexes with each other was evaluated. FLAG-JB12 was introduced into cells that were challenged with P23H-R or P23H-C110S-C187S-R. Both P23H-R and P23H-C110S-C187S-R were present in FLAG-JB12 immunoprecipitates with Hsp70, but the ER-transmembrane chaperone calnexin was absent (Figure 2C).

To test the functional importance of JB12’s interaction with Hsp70 in triage events, P23H-R was co-expressed with either WT JB12 or QPD-JB12. QPD-JB12 is a dominant-negative mutant that binds non-native clients and interacts with protein homeostatic machinery, but its Hsp70 binding domain is defective due to a Q to H mutation in its HPD-motif. QPD-JB12 is incapable of functional interactions with Hsp70, so it binds non-native clients and blocks their down-stream binding to Hsp70 (Grove et al., 2011; He et al., 2021). Western blot analysis revealed a greater than 4-fold increase in P23H-R steady state levels in the presence of QPD-JB12, whereas elevation of WT-JB12 reduced steady-state levels of P23H-R in a dose dependent manner (Figure 2D). Notably, QPD-JB12 associates with P23H-R at higher levels than WT-JB12 (Figure 2E) and does not cause P23H-R to enter into detergent insoluble aggregates (Figure S2B and C). Imaging of cells co-expressing P23H-R and QPD-JB12 showed reduced colocalization of P23H-R with lysosomes and retention in the ER (Figure S2C). Expression of dominant-negative mutant forms of the core autophagy initiation kinase ULK1 (Unc-51 like autophagy activating kinase) or Beclin-1, the scaffold for the Class III PtdIns3*P* kinase VPS34, prevented P23H-R accumulation in lysosomes with LAMP (Figure S2C). Additionally, treating cells with the VPS34 inhibitor PIK-III caused a significant decrease in P23H-R colocalization with LAMP1 (Figure S2D). JB12 and Hsp70 are present in complexes with P23H-R and function in a pathway requiring the canonical autophagy initiation factors ULK1, and Beclin-1 to triage P23H-R intermediates for degradation via ERAA.

In unstarved cells, FIP200 (focal adhesion kinase family interacting protein of 200 kD) localizes on intracellular membranes to target its partner, ULK1, to focally initiate selective-autophagy (Ravenhill et al., 2019; Vargas et al., 2019). Hsp70 and Hsp40s cooperate with Hsp90 to regulate the activity of kinases that include ULK1 (Joo et al., 2011). It is therefore possible that interaction of JB12 with FIP200 could determine sites in the ER where ERAA occurs and/or regulate client flux through the pathway. To address this possibility we conducted FLAG-JB12 immunoprecipitations from extracts of cells expressing P23H-R. FIP200 and Hsp70 were present in the products of FLAG-JB12 pull-downs. Levels of FIP200 and P23H-R were elevated in immunoprecipitates with FLAG-QPD-JB12, but Hsp70 was absent (Figure 2E). Thus, JB12 is detected in association with FIP200, Hsp70, and P23H-R. QPD-JB12 might inhibit ERAA because it sequesters FIP200 in complexes that cannot be activated due to the absence of Hsp70. The association of FIP200 in complexes with JB12 and P23H-R suggests a potential role for JB12 in local regulation of FIP200/ULK1 function in ER-microdomains where ERAD-RMPs accumulate. These data support the concept that JB12 participates in the focal regulation of ERAD-RMP flux through ERAA (He et al., 2021; Houck et al., 2014).

### Mechanisms for JB12-Dependent Triage of P23H-Rhodopsin are Distinct from ER-Phagy

When the ER is challenged with starvation, chemical damage, or protein aggregate accumulation there are several mechanisms for selective-ER-phagy of damaged microdomains (Chino and Mizushima, 2020; Ferro-Novick et al., 2021; Loi and Molinari, 2020; Wilkinson, 2019b). We propose that JB12 and Hsp70 act to clear ERAD-RMPs via selective-autophagy to protect the ER from damage in healthy biosynthetically active cells to suppress the induction of ER-phagy. To further understand this process, we uncovered the route for passage of P23H-R from the ER to lysosomes. At the onset, we compared the localization of P23H-R in lysosomes to the location of ER-proteins that would be degraded by ER-phagy (Figure 3A-F). P23H-R accumulates predominantly in lysosomal membranes of live-cells with LAMP1-mCherry even after the addition of BafA1 (Figure 3A). To corroborate this observation at a higher resolution, transmission electron microscopy (TEM) was conducted with cells expressing P23H-R and immunogold stained for P23H-R. P23H-R is detected in the lysosomal membrane, which is distinct from ER-whorls present in the lumen (Figure 3B). The presence of P23H-R within lysosomal membranes is not typical for clients of selective or basal autophagy and supports a model in which ERAD-RMPs are sorted within the ER-tubular network directly into the membranes of ER-connected phagophores (Hayashi-Nishino et al., 2009; He et al., 2021; Houck et al., 2014; Yla-Anttila et al., 2009).

**Figure 3.**
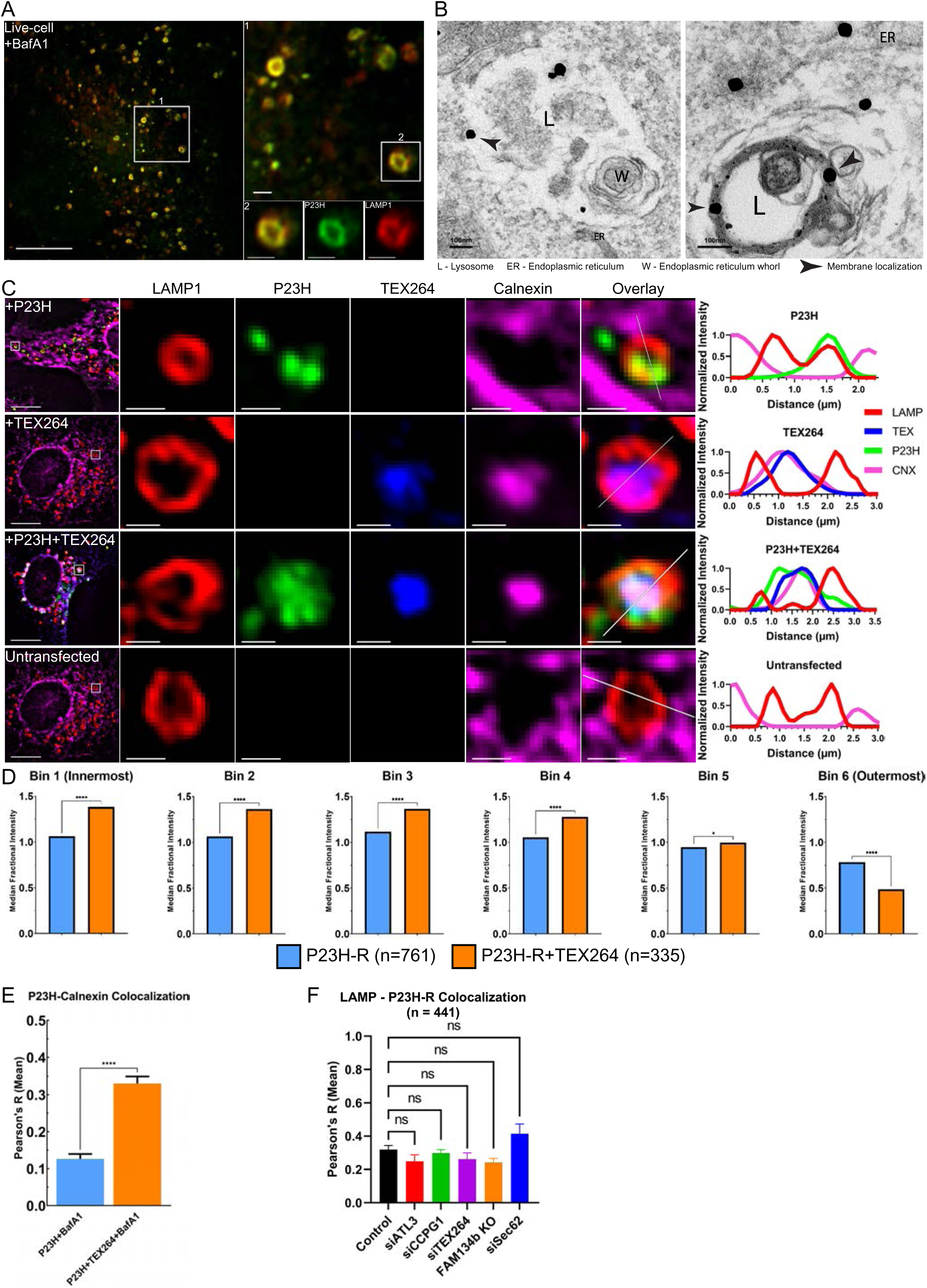
P23H-Rhodopsin is trafficked independent of ER-phagy indicators. (A) Live-cell confocal microscopy of on Zeiss 880 with Airyscan of COS-7 cells transiently transfected with P23H-R and LAMP1-mCherry (B) TEM image of anti-GFP immunogold staining in COS-7 cells transiently transfected with P23H-R. Scale bar = 100nm (C) Left: Fixed-cell epifluorescent microscopy: COS-7 cells were transiently transfected with either P23H-R (Green), TEX-164-HaloTag (Blue) or both, then incubated with 50nM Janelia Fluor^®^ 646 HaloTag^®^ Ligand (blue) and 100nM BafilomycinA1, followed by fixation and immunofluorescence staining for LAMP1 (Red) or Calnexin (Magenta). Right: Plot profiles for line drawn in inset overlay (D) Median fractional intensity of P23H-R for each segment within identified LAMP1 objects; Mann-Whitney test. Bin 1 corresponds to the innermost segment of the LAMP1 object (i.e. center point inside LAMP1 ring), whereas Bin 6 is the outermost (i.e. outer rim of LAMP1 ring). Mann-Whitney test (E) Bar graph of mean Pearson’s R values between Rhodopsin and Calnexin calculated within identified LAMP1 objects from Figure 3C. Mann-Whitney test and (n=1096; error bars for SEM) (F) Mean of Pearson’s R calculated across multiple fields of cells for each indicated condition (error bars for SEM). All ER-phagy receptors were knocked down by siRNA, except for FAM134b which used a CRISPR knockout cell line generated via lentiviral transduction. Kruskal-Wallis and Dunn’s multiple comparisons tests \**P* < 0.05, \*\*\*\**P* < 0.0001 Scale bars (unless indicated): whole cell views = 10µm; insets = 1µm

If this model is accurate, then P23H-R localization in the lysosome will vary according to the pathway; in ERAA, P23H-R resides in the lysosome membrane, while in ER-phagy and as a casualty of ER fragmentation P23H-R should be located in the lumen. TEX264 facilitates the degradation of damaged ER tubules that are engulfed by phagophores and delivered to the lysosome lumen via fusion of the autophagosome outermembrane with the lysosome membrane (An et al., 2019; Chino et al., 2019; Delorme-Axford et al., 2019). The localization of P23H-R was therefore imaged along with endogenous calnexin and LAMP in control cells, and also in TEX264-HaloTag overexpressing cells. P23H-R was detected in the membrane of LAMP-positive lysosomes that contact the ER, but are devoid of calnexin. In cells expressing exogenous TEX264-HaloTag, TEX264 and calnexin accumulated within the lysosomal boundaries LAMP, indicating the occurrence of ER-phagy. When exogenous TEX264 was co-expressed with P23H-R, ER-phagy shifted the location of P23H-R to the lumen with calnexin and TEX264 (Fig 3C-E).

To verify this observation was consistent across vesicles and cells, LAMP vesicles were identified as in Figure 1D. This was followed by segmenting each vesicle with equidistant concentric circles, with each annuli serving as a bin for the total intensity of P23H-R observed within the object. Where bin 1 corresponded to the center of the lysosome, and Bin 6 to the outer edge of the lysosomal membrane, fractional intensities were calculated across all bins (fraction of total object intensity within each bin). In counts of 1096 vesicles, statistically significant differences between ‘P23H-R’ and ‘P23H-R+TEX264’ were seen across all 6 bins, confirming that increases in TEX264 activity shifts the localization of P23H-R from the lysosome membrane to lumen. Sorting of P23H-R to the lysosome membrane appears selective because calnexin is not normally present in lysosomes, whereas when TEX264 is overexpressed, calnexin is detected in the lysosome lumen with P23H-R.

To evaluate the potential role of ER-phagy receptors in selective P23H-R triage, we used CRISPR/Cas-9 and siRNA to deplete each ER-phagy candidate from cells (Chino and Mizushima, 2020; Ferro-Novick et al., 2021; Loi and Molinari, 2020; Wilkinson, 2019b) (Figure 3F and Figure S3). The impact that loss of each respective ER-phagy receptor had on the co-localization between P23H-R and LAMP in fixed cells was evaluated by Pearson’s correlation across 441 cells. Each respective knock-down cell line failed to reveal a statistically significant difference, showing that depletion of every known ER-phagy receptor did not affect P23H-R accumulation in lysosomes (Figure 3F and Figure S3). These observations strongly suggest an ER-exit mechanism for selective-autophagy of P23H-R that distinct from events that involve the engulfment by phagophores of ER-membranes containing a particular ER-Phagy receptor. This process requires JB12 and Hsp70, is dependent upon the conformation of misfolded membrane protein intermediates, employees the canonical autophagy initiation machinery ULK1 and Beclin-1, and is selective because it does not consume resident ER-membrane chaperones such as calnexin and JB12 or globally misfolded membrane protein intermediates.

### P23H-R Intermediates Pass Through Omegasomes During Passage to Lysosomes

ERAD-RMPs are associated with JB12 and Beclin-1/VPS34 complexes during the initiation of ERAA, and these events are proposed to provide a mechanism for focal activation of ERAA in biosynthetically active ER-microdomains (He et al., 2021; Houck et al., 2014). This helps explain why ER-phagy receptors are dispensable for degradation of ERAD-RMPs. A key feature of this model that would explain the presence of P23H-R in lysosome membranes is that ERAD-RMP containing ER tubules are the source of the phagophores for ERAA.

Consistent with the concept that JB12 facilitates the client-dependent flux of ERAD-RMPs through ERAA, we detect the localization of endogenous JB12 in discrete foci with endogenous FIP200 at the edge of a larger area enriched in P23H-R (Figure 4A). The co-localization of endogenous JB12, FIP200, and P23H-R is consistent with the presence of endogenous FIP200 and P23H-R in the products of immunoprecipitants with FLAG-JB12 and the requirement for ULK1 in ERAA (Figure 2 and Figure S2).

**Figure 4.**
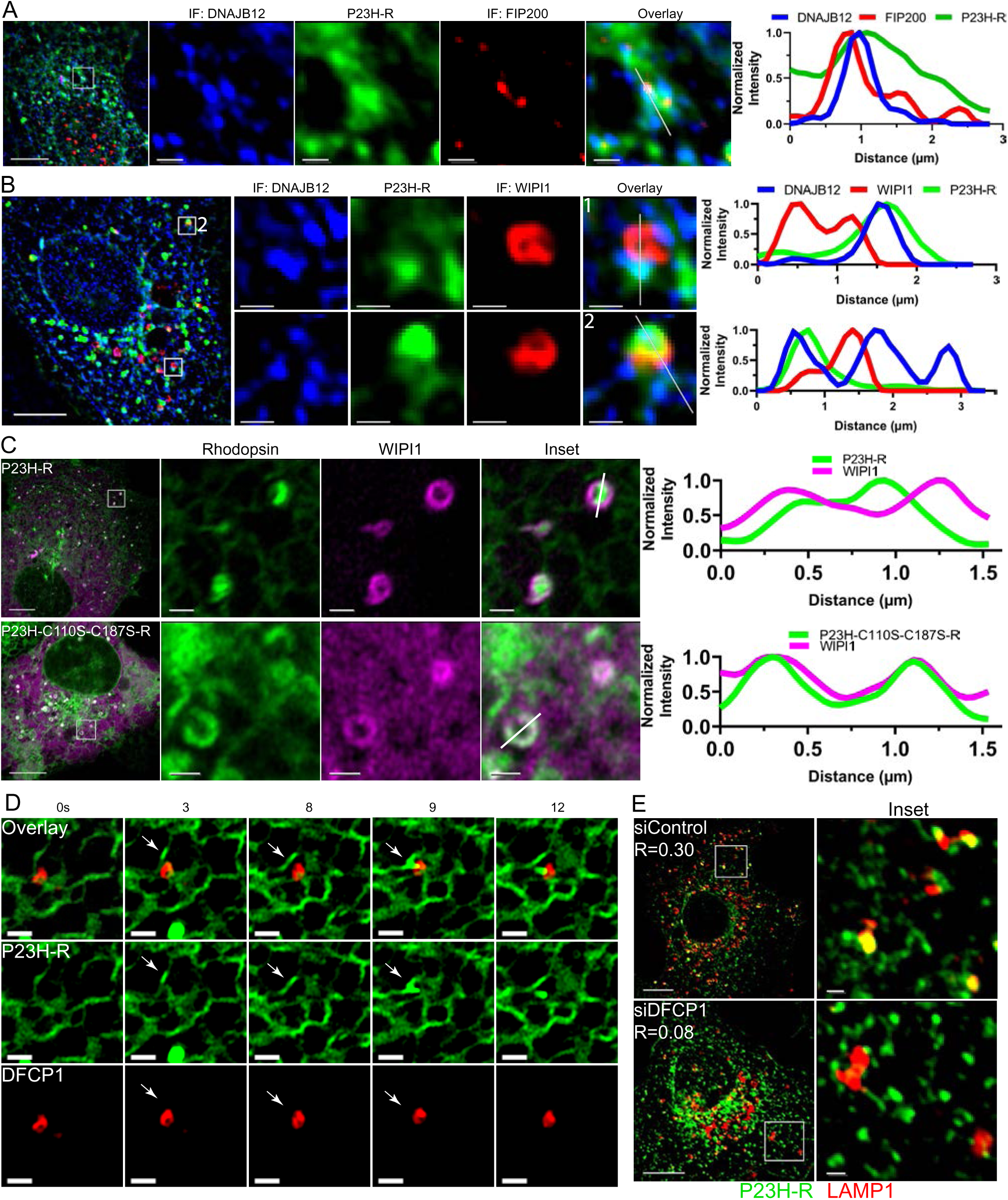
P23H-Rhodopsin Localizes to Omegasome Markers. (A-B) Left: Fixed-cell epifluorescent microscopy of COS-7 cells transiently transfected with P23H-R and immunostained for indicated marker. Right: Plot profile for line drawn in inset overlays (C) Left: Live-cell confocal microscopy of COS-7 cells expressing WIPI1-mCherry and either P23H-R or P23H-C110S-C187S-R. Right: Plot profile of line drawn in inset overlay (D) Time-lapse of live-cell confocal microscopy of COS-7 cells expressing DFCP1-mCherry and P23H-R (E) Fixed-cell epifluorescent microscopy of COS-7 cells expressing P23H-R after siControl or siDFCP1 knockdown

Foci of different sizes that are enriched in endogenous JB12 and P23H-R, colocalized on the edge of omegasomes marked by endogenous WIPI1 rings (Figure 4B, inset 1 and inset 2). In live-cell imaging where WIPI1-mCherry was co-expressed with either P23H-R or P23H-C110S-C187S-R, P23H-R signal was found inside WIPI1-mCherry rings in membranes that remained connected to the ER (Figure 4C). In contrast, P23H-C110S-C187S-R did not enter inside WIPI1 rings, but did colocalize with WIPI1 (Figure 4C). This colocalization is expected since P23H-C110S-C187S-R is present in ER-membranes and some of these are painted with endogenous WIPI1, but P23H-C110S-C187S-R does not enter ERAA.

DFCP1 acts in concert with WIPI1 during omegasome formation (Tooze and Yoshimori, 2010). In live cells, DFCP1-rings were of similar size as WIPI1 rings and contained P23H-R enriched ER-membranes (Figure S4B). At t=3 and 8s, an ER tubule containing P23H-R is seen adjacent to a DFCP1 ring. At t=9 seconds a portion of the tubule contacts the DFCP1-ring. Thus, ER tubules containing P23H-R dynamically interact with omegasomes.

There are four WIPI family members in primate cells (De Leo et al., 2021; Thurston et al., 2016) and we were not able to deplete all members of the WIPI family to levels that block basal-autophagy or ERAA (not shown). There are only two DFCP homologs expressed in mammals and siRNA depletion of DFCP1 was sufficient to impair the accumulation of P23H-R in lysosomes (Figure 4E). Therefore, P23H-R-enriched ER tubules dynamically associate with components of omegasomes that are necessary for its accumulation in lysosomes. These data support a model in which ER tubules containing P23H-R are converted into phagophores resultant from JB12s interaction with ERAD-RMPs and components of the autophagy initiation kinase complexes.

### Lysosomes Dynamically Dock with WIPI1 rings Containing P23H-Rhodopsin Enriched ER tubules

Next, we used fixed and live-cell imaging to study the mechanisms for transfer of P23H-R from the ER to lysosomes (Figure 5). We stained fixed cells for endogenous WIPI1 and LAMP and sought to detect P23H-R-enriched autophagosomes docking to lysosomes that are located in the vicinity of the centrosome (Pu et al., 2016). Instead, we were surprised to observe the association of lysosomes with WIPI1 rings (Figure 5A). These ‘docking events’ were transient, yet stable, with the longest association seen to last 70 seconds (Figure S5A). Object-based image analysis was used to evaluate the association of LAMP with WIPI1 foci, which showed 10% of the 9823 identified WIPI1 objects as overlapping with LAMP. However, in cells expressing P23H-R, 25% of the 743 WIPI1 objects overlapped with LAMP. Thus, lysosomes dock to WIPI1 rings and may do so to accept autophagic cargo that emerge from local phagophores. This docking reaction is stimulated to a 250% increase when challenged by P23H-R, which is consistent with lysosome docking to omegasomes being important for ERAA.

**Figure 5.**
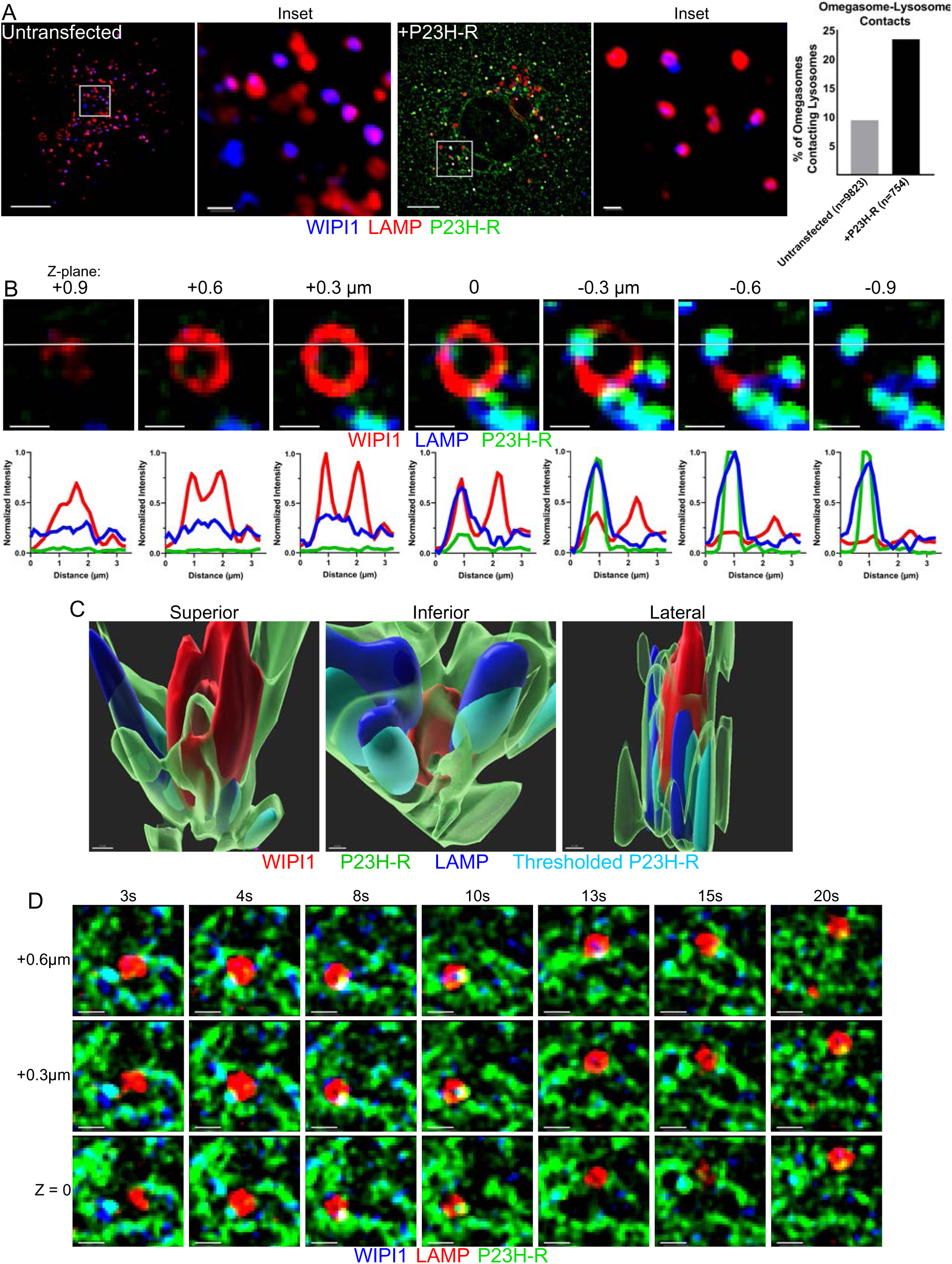
Live-cell Imaging Reveals P23H-Rhodopsin Transfer to Lysosomes occurs at Omegasome Contacts. (A) Left: Fixed-cell epifluorescent microscopy of COS-7 cells expressing P23H-R (Green) with immunofluorescence staining for LAMP1 (Blue) or WIPI1 (Red). Inset for ‘+P23H-R’ condition shows muted P23H-R channel. Right: Bar graph of LAMP1-WIPI1 contact frequencies seen in each condition from object detection. Frequency was calculated by dividing number of LAMP1-masked omegasome objects by the total number of omegasomes identified. Scale bars: whole cell views = 10µm; insets = 1µm Live-cell Confocal Microscopy with resonant scanning of COS-7 cells expressing P23H-R, WIPI1-mCherry, and mTagBFP2-LAMP1 (B) Slices of Z-stack with plot profiles of drawn line underneath each respective image. Scale bar = 1µm (C) 3D view of surface renders for the Z-stack shown in *(B)*. ‘Thresholded P23H-R’ refers similar intensity thresholding (top 10% of histogram intensities) as displayed in *(B)*. ‘P23H-R’ in transparent green includes the top 90% of intensities recorded for P23H-R in the Z-stack. Scale bar = 0.1µm (D) Additional Z-stack time-lapse of mTagBFP2-LAMP1 docking and departing from WIPI1-mCherry site containing P23H-R

Images of live-cells expressing WIPI1-mCherry, LAMP1-mTagBFP2, and P23H-R permitted the capture of a 3-dimensional view of the interface between WIPI1 rings, P23H-R phagophores, and lysosomes. Examination of a Z-stack containing the WIPI1 ring and a docked lysosome suggests a potential mechanism by which P23H-R exits the ER during ERAA. Stepwise slices of the Z-stack that are +0.9 to −0.9 μm above or below the center of the WIPI1 structure show that P23H-R is enriched in lysosomes that dock with one side of the WIPI1 ring at −0.3 to −0.9 μm (Figure 5B). 3-Dimensional views of this Z-stack processed are presented as superior, inferior, and lateral views (Figure 5C and Figure S5B). The superior view displays WIPI1 in the shape of a cylinder that is contacted along its exterior surface by P23H-R enriched ER tubules. There is also a P23H-R enriched ER-membrane that coats the wall of the WIPI1 ring, then folds into its central cavity, and then out the top. The inferior view displays a notch in the WIPI1 ring in which a lysosome has docked. The lateral view shows P23H-R that co-localizes with LAMP1 in lysosomes. We interpret these data to suggest that P23H-R enriched ER tubules interact with omegasomal surfaces containing autophagy initiation machinery. These interactions then facilitate the metamorphosis of P23H-R tubules into phagophores that deliver their contents to lysosomes docked on the same WIPI1 ring.

To test this model, the dynamic association of lysosomes with WIPI1 rings containing P23H-R-tubules was evaluated in a live z-stack acquisition over time (Figure 5C and Figure S5). Shown are images at z=0, z=0.3 and z=0.6μm between t=3s to t=20s. A lysosome is seen separate, but nearby, an omegasome containing WIPI1 and P23H-R (t=3s), followed by its approach (t=4s), then docking (t=8 to 10s) to the omegasome. The lysosome disengages (t=13s), moves away from the WIPI1 ring (t=15s), and now contains P23H-R in its membrane (t=20s). These collective data demonstrate the dynamic association of lysosomes with WIPI1 rings that contain an ERAA client. They support the interpretation that autophagic transfer of P23H-R out of the ER occurs during lysosome docking to WIPI1 rings.

### GABARAP Facilitates P23H-R Exit from Omegasomes and Accumulation in Lysosomes

The ER tubules containing P23H-R extend from omegasome-lysosome contacts, and we surmise these tubules are of ER-connected phagophores (Hayashi-Nishino et al., 2009, 2010; Yla-Anttila et al., 2009). We therefore evaluated the requirement of the ATG8 family members LC3a/b/c and GABARAP/L1/L2 in clearance of P23H-R from the ER. The mammalian ATG8 family members LC3a/b/c and GABARAP/L1/L2 contain binding sites for microtubules, and are conjugated through their C-termini to the head groups of phosphatidyl-ethanolamine present in omegasomes at the onset of phagophore biogenesis (Johansen and Lamark, 2019). ATG8s also contain a domain that recognizes LIR-motifs (LC3 interacting region) present in autophagic cargo-receptors and also autophagy initiation machinery such as FIP200 and ULK1 (Johansen and Lamark, 2019). To initiate this investigation, the localization of signals from 3xFLAG-GABARAP and LC3b relative to P23H-R in a representative lysosome was analyzed. Signals for P23H-R and 3xFLAG-GABARAP co-localized in the lysosome membrane (R=0.633), with markedly lower P23H-R:LC3B colocalization (R=0.094) since the signal for LC3B is lumenal (Figure S6A).

Unbiased object-based quantitation of over 13,700 lysosomes showed that the simultaneous depletion of GABARAP, GABARAP-L1, and GABARAP-L2 by siRNA (siGABARAP/L1/L2) decreased the accumulation of P23H-R in lysosomes (Figure 6A and Figure S6G). Importantly, siGABARAP/L1/L2 did not observably impede autophagic flux since it did not abolish LC3b trafficking to lysosomes (Figure S6B). Due to functional redundancy, when GABARAP, GABARAPL1, and GABARAPL2 were knocked down individually, a statistically significant decrease in accumulation of P23H-R foci with LAMP was not observed (not shown) (Vaites et al., 2018). These observations were verified in stable HeLa cells possessing complete knockout of the ATG8 families by CRISPR/Cas9 (Figure S6L); GABARAP/L1/L2 KO impaired P23H-R accumulation in lysosomes, yet this trafficking was restored after co-transfection with GABARAP (Figure S6K). Both the complete knockout and siRNA knockdown of the LC3 subfamily was not observed to impede P23H-R localization to lysosomes. Additionally, the autophagic cargo receptor p62 is a ubiquitin-binding autophagy receptor for misfolded proteins and protein aggregates, but it localizes to the center of lysosomes, and not with P23H-R in the lysosomal membrane (Figure S6C).

**Figure 6.**
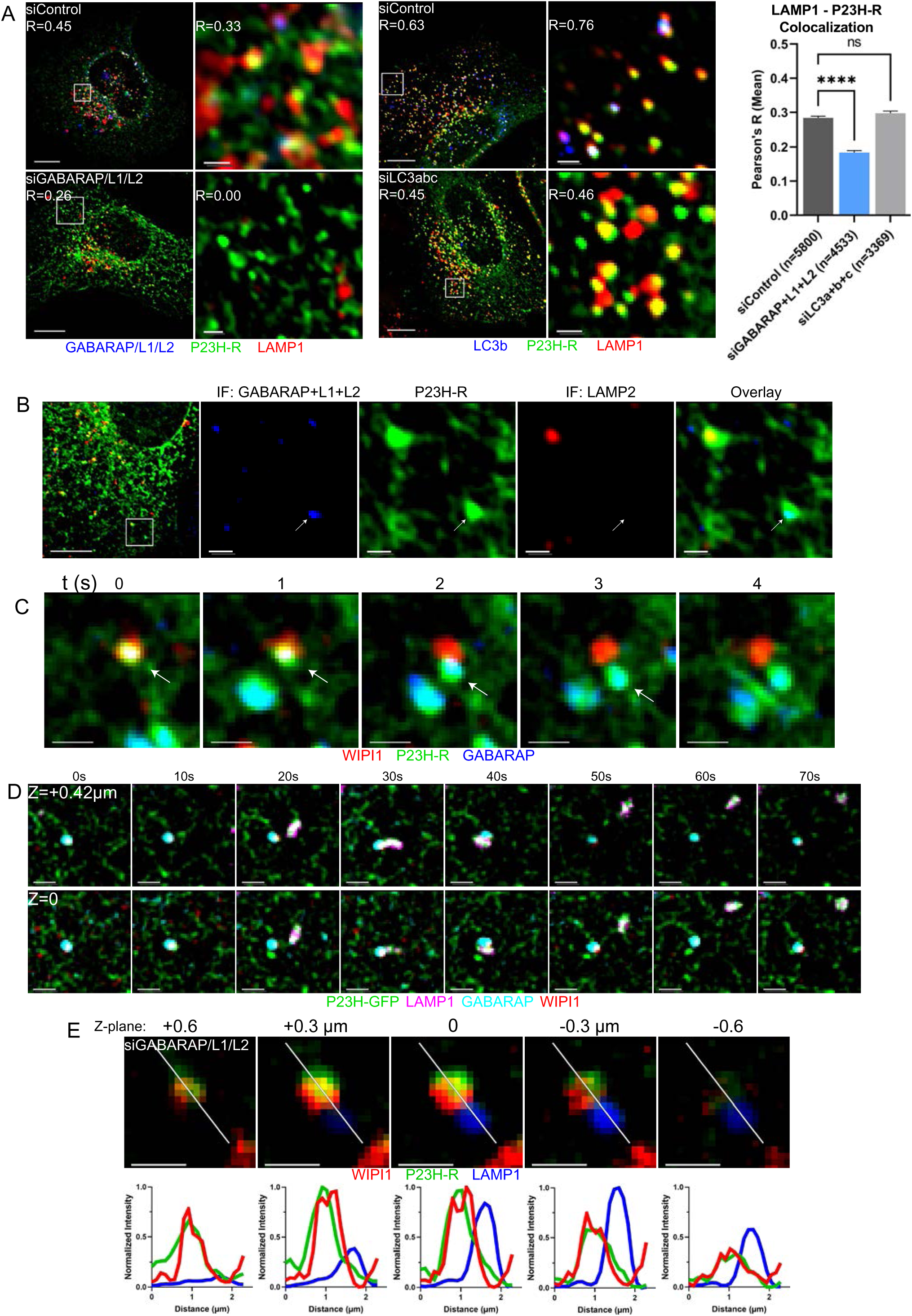
P23H-Rhodopsin Associates with Specific Autophagic Cargo Receptors. (A) Left: Fixed-cell epifluorescent microscopy of COS-7 cells expressing P23H-R after siControl, siGABARAP/L1/L2, or siLC3abc knockdown. Right: Bar graph of mean Pearson’s R values between Rhodopsin and LAMP1 calculated within identified LAMP1 objects. Kruskal-Wallis and Dunn’s multiple comparisons tests (B) Fixed-cell epifluorescent microscopy of COS-7 cells expressing P23H-R and immunolabeled for LAMP2a and GABARAP/L1/L2 (C) Time lapse from live-cell confocal microscopy with resonant scanning of COS-7 cells expressing P23H-R, WIPI1-mCherry, and HaloTag-GABARAP and incubated with 50nM of Janelia Fluor^®^ 646 HaloTag^®^ Ligand for 30min before imaging (D) Same as in *6D* but also expressing mTagBFP2-LAMP1. Manual X offset correction for P23H-R and HaloTag-GABARAP = +11pixels (E) Live-cell confocal microscopy with resonant scanning of COS-7 cells expressing P23H-R, WIPI1-mCherry, and mTagBFP2-LAMP1. Slices of Z-stack with plot profiles of drawn line underneath each respective image. Scale bar = 1µm ‘GABARAP/L1/L2’ includes GABARAP, siGABARAP-L1, siGABARAP-L2 concurrently; ‘LC3abc’ includes LC3a, LC3b, and LC3c concurrently Scale bars: whole cell views = 10µm; insets = 1µm

The punctuate nature of the P23H-R and GABARAP signal in the lysosome membrane suggests that P23H-R and GABARAPs move in the same compartment from the ER to lysosomes (Figure S6A). If this supposition is accurate then GABARAPs should localize in phagophores with P23H-R. Images obtained in fixed COS-7 cells identify ER-microdomains that are enriched with GABARAP and P23H-R that do not contain LAMP (Figure 6B). The presence of GABARAP with P23H-R, departing together from WIPI1 rings, was demonstrated via 3-color live-cell imaging in cells expressing WIPI1-mCherry, HaloTag-GABARAP, and P23H-R (Figure 6C). GABARAP and P23H-R are present in phagophore membranes that extend into the cytosol from WIPI1-decorated omegasomes.

To monitor the localization of P23H-R and GABARAP within the phagophore membrane and the movement of P23H-R to lysosomes containing GABARAP, we conducted 4-color live-cell imaging for cells expressing P23H-R (green), WIPI1-mCherry (red), HaloTag-GABARAP (cyan), and LAMP1-mTagBFP2 (magenta) (Figure 6D and S6D). A lysosome (already containing P23H-R) was observed to dock (t=20 to 40s) at an ER-microdomain containing WIPI1, P23H-R, and GABARAP, followed by the lysosome’s departure (t=50s). The WIPI1 foci remained intact after lysosome departure. These data demonstrate the presence of GABARAP in phagophores with P23H-R and also in lysosomes that contain P23H-R transiently associating with WIPI1 rings.

Analysis of docking times between WIPI1 and LAMP1 varied from short interactions (less than 10s), to associations that lasted up to 70s (Figure S5A), with the average docked time at 22 seconds (Figure S6F). siGABARAP/L1/L2 was not observed to interfere with the WIPI1:lysosome docking, nor change the average docking time of LAMP1 to omegasomes (Figure S6F). Therefore, docking of lysosomes to WIPI1 foci does not appear to require GABARAP.

A close view of a WIPI1 ring containing a docked lysosome in live siGABARAP/L1/L2 cells shows the presence of P23H-R phagophores extending from the central region of a WIPI1 ring. There is a lysosome docked to the WIPI1-decorated ER surface containing a P23H-R enriched phagophore, however the docked lysosomes do not contain P23H-R (Figure 6E). This mirrors what was observed in GABARAP/L1/L2 knockout cells, as ER-connected P23H-R foci were seen abutting lysosomes (Figure S6K). These results are in contrast to normal cells where P23H-R and LAMP signals overlap at the edge of a WIPI1 ring (Figure 5B). Terminal autophagic transfer of P23H-R from phagophores to lysosomes appears to require GABARAP.

To understand why GABARAP/L1/L2, but not LC3a/b/c depletion, blocks P23H-R exit from omegasomes, we added purified GST-GABARAP and GST-LC3b to HEK293 cell extracts and performed pull-downs (Figure S6H and S6I). GST-GABARAP and GST-LC3b pulled down JB12 and the ER-phagy receptor CCPG1 and TEX264. Remarkably, GST-GABARAP, but not GST-LC3b, pulled down FIP200, and ULK1. The JB12 homolog JB14, which is not required for ERAA, was not present in pulldowns. Thus, JB12, FIP200, and ULK1 are present in GABARAP-containing complexes and ULK1 absent for LC3 containing complexes. In addition, FIP200, ULK1, and JB12 co-eluted with GABARAP in a pull-down of cells expressing 3xFLAG-GABARAP (Figure S6J). These data are consistent with previous observations on the specificity of interactions between different GABARAP and LC3 family members with FIP200 and ULK1 (Grunwald et al., 2020). Endogenous JB12 and FIP200 co-localize with P23H-R in small foci within omegasome rings (Figure 4A) and JB12, ULK1 and FIP200 are present in complexes with GABARAP (Figure S6H and S6J). The collective data presented suggest a model in which JB12 delivers P23H-R to omegasomes, and that GABARAP facilitates P23H-R movement from the ER to lysosomes that dock with WIPI1 rings (Figure 7).

**Figure 7.**
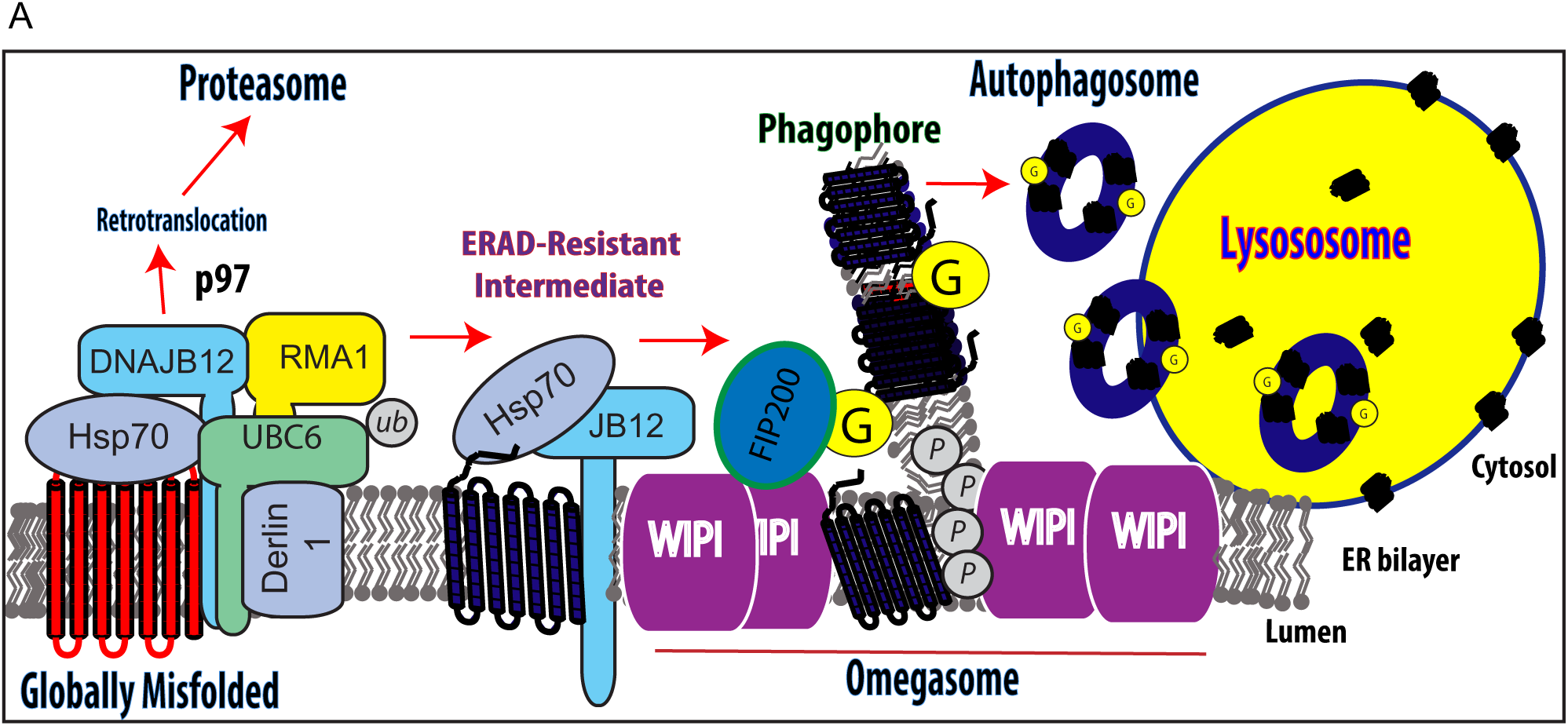
Model for the Degradation of Misfolded Membrane Proteins by ERAA. (A) Illustration of the model describing the triage of ERAD-resistant membrane proteins to lysosomes

## Discussion

Study of the mechanisms for triage of misfolded rhodopsin degradation intermediates has revealed points of cross-talk between the Hsp40/Hsp70 chaperone system and canonical autophagy initiation machinery during lysosomal degradation of ERAD-RMPs (Figure 7). Globally misfolded P23H-R is degraded by ERAD, whereas biochemically stable and non-aggregated P23H-R intermediates are cleared from the ER by ERAA. ER-transmembrane JB12 and Hsp70, which act as substrate selectors for ERQC-E3 ligases to mediate ERAD of globally misfolded membrane proteins, facilitate both ERAD and ERAA of P23H-R (Grove et al., 2011; Houck et al., 2014; Sopha et al., 2017). ERAD-RMPs interact with JB12/Hsp70 complexes for extended time-periods, and JB12 is necessary for entry of P23H-R into the ERAA pathway. JB12 and FIP200 co-localize with P23H-R in foci on the rim of omegasomal rings and are present in complexes with GABARAP. Hsp70 and Hsp90 regulate the activity of kinases that include ULK1, so JB12 and Hsp70 interactions with FIP200 and ULK1 could be of a regulatory nature(Joo et al., 2011). In such a scenario, JB12 would have dual functions: one - as a molecular chaperone that maintains P23H-R in a degradable detergent soluble state; two - in the focal regulation of ULK1/FIP200 action on ERAD-RMP containing ER tubules that enter or associate with the surface of omegasomes.

Data presented identifies networks of interacting ER tubules that contain or are decorated with different cellular components that facilitate the clearance of P23H-R from the ER. One set of ER tubules contains JB12 and P23H-R, and we detect them in association with a separate network of ER-membranes that are coated with DFCP1 and WIPI1. WIPI1 rings contain the autophagy initiation machinery that facilitates phagophore formation (Joachim and Tooze, 2016; Ktistakis and Tooze, 2016). Discrete foci containing JB12, FIP200, and P23H-R decorate the edge of omegasomes, with adjacent P23H-R tubules threading along the walls of WIPI1 rings. These tubules are devoid of JB12, yet contain GABARAP and represent ER-connected phagophores. These data are consistent with previous reports of ER-connected phagophores that were detected by TEM(Hayashi-Nishino et al., 2009), and are a feature of ERAA that is distinct from ER-phagy.

The presence of P23H-R in ER-connected phagophores permits their analysis in live-cells, and data reported suggest that entrance of ERAD-RMPs into them is selective. The ER-resident chaperones JB12 and calnexin are excluded from these phagophores, as are globally misfolded membrane proteins. JB12 is required for ERAD-RMP sorting into ER-connected phagophores, but the mechanism for selective entry or retention of ERAD-RMPs within them is not yet clear. Nevertheless, Hsp70 and Hsp40s can sequester misfolded proteins into soluble and non-toxic oligomers, so JB12 and Hsp70 have the capability to organize ERAD-RMPs into assembles that could be selectively sequestered into ER-connected phagophores (Ho et al., 2019; Wolfe et al., 2013).

A novel feature of ERAA is that the transfer of ERAD-RMPs to lysosomes is associated with lysosome docking to WIPI1 rings. The association of lysosomes with WIPI1 rings occurs over 10-70 second time spans. In control cells around 10% of all endogenous WIPI1 foci show a docked lysosome, and that docking frequency increases 2.5-fold upon P23H-R expression. Docking of lysosomes to WIPI1 rings does not require GABARAP, but the transfer of P23H-R from the ER to lysosomes is GABARAP-dependent. 3-dimensional reconstruction of images in Z-stacks taken in live-cells of a WIPI1 ring, P23H-R phagophore, and lysosome, show the extent to which these different structures are in contact. Lysosomes are observed to dock in a notch on the distal end of a WIPI1 ring and are enriched in P23H-R. ERAD-RMP containing ER-membranes also envelope WIPI1-associated lysosomes which provides an interface where efficient transfer of P23H-R from the ER to lysosomes can be coupled to docking events.

Lysosomes dynamically interacting with omegasomes containing P23H-R phagophores was unanticipated. Lysosomes are highly dynamic organelles that can dock with the ER where the presence of PtdIns3*P* on lysosomes is recognized by ORP1L and ER-localized protrudin in a process that is regulated by the PtdIns3*P* kinase VPS34 (Pedersen et al., 2020). If ORP1L and protrudin were present in omegasomes, they could mediate the observed lysosome docking to them. Furthermore, WIPI1 was recently shown to oligomerize and insert its amphipathic helices into tubular membranes of endocytic organelles to drive their fission (De Leo et al., 2021). It is therefore possible that WIPI1 mediates the fission of P23H-phagophores membranes during P23H-autophagosome formation in a process that is coordinated with lysosome docking. Lysosome docking and cargo transfer also describes chaperone-mediated autophagy (CMA), a process mediated by the Hsp70, HSPA8, whereby cargo are directly transferred across lysosome membrane. However CMA is noted as a VPS34-independent process (Klionsky et al., 2021), which excludes ERAA as our data show involvement of ULK1, Beclin-1 and VPS34 (Figure 2 supplement).

Depletion of GABARAP by siGABARAP/L1/L2 leads to the arrest of P23H-R autophagic degradation after it has accumulated in phagophores, but prior to P23H-R transfer to adjacent lysosomes. GABARAP is therefore not required for the formation of P23H-R phagophores. A mechanism for GABARAP function in P23H-R transfer from phagophores to lysosomes is suggested by the observation that GABARAP, FIP200, and ULK1 are present in the same complexes. The GABARAP present in omegasomes could therefore mediate localization of ULK1 to locations where its action is required to induce liberation of autophagosomes containing P23H-R. Such a regulatory mechanism would enable a two-step protection of the ER from toxic misfolded membrane proteins. In step 1, sequestration of P23H-R into phagophores occurs to insulate a dominantly toxic P23H-R intermediate away from other ER-components. Step 2 would involve the transfer of P23H-R out of the ER in reactions catalyzed by GABARAP that are coordinated with lysosome docking to WIPI1 rings.

The triage of biochemically stable intermediates of P23H-R to ERAA identifies a cellular mechanism to prevent ERAD-RMPs from saturating JB12 and causing ER-stress induced apoptosis (Sopha et al., 2017). JB12 functions with the ERQC E3-ubiquitin ligase GP78 to facilitate the constitutive degradation of the short-lived and ER-associated BCL-2 homolog BOK (Llambi et al., 2016; Sopha et al., 2017). ER-stress dependent degradation of JB12 causes BOK accumulation and primes cells for apoptosis (Sopha et al., 2017). Data presented and those from related studies suggest that JB12 and Hsp70 maintain ERAD-RMPs in a detergent soluble and non-aggregated state that is mobile in the ER-membrane (He et al., 2021). JB12 is not stress inducible, so cells clear ERAD-RMPs from it by creating ER-tubule networks that feed ERAD-RMPs into ER-connected phagophores formed in association with WIPI1 rings. JB12 is present in complexes with ERAD-RMPs, FIP200, ULK1, and GABARAP, yet JB12 is not detected in lysosomes with P23H-R and does not appear to be consumed during ERAA. Thus, JB12 appears to sense ERAD-RMP accumulation and work with Hsp70 to facilitate the focal activation of ERAA by associating with complexes containing FIP200 in omegasomes. This mechanism prevents JB12 from forming dead-end complexes with ERAD-RMPs to permit the constitutive degradation of BOK.

## Acknowledgments

This work was supported by grants from the NIH (GM056981). The UNC Hooker and Neuroscience Imaging Core housed the Olympus FV3000 microscope. Dr. Michael Cheetham, University College of London, England, kindly provided the rhodopsin expression plasmids. Transmission electron microscopy was performed at the UNC Department of Pathology Microscopy core. We also thank Wade Harper, Harvard University, for generously providing the ATG8 knockout cell lines.

## Materials and Methods

### Reagents

DC Protein Assay (500-0119) and Clarity Western ECL Substrate (170-5061) from BioRad. Effectene Transfection Reagent (301427) from Qiagen. Lipofectamine RNAiMAX Reagent (100014472) from Thermo Fisher Scientific. ProLong Diamond antifade mountant without (P36970) or with DAPI (P36971) from Thermo Fisher Scientific. Bafilomycin A1 (11038) from Cayman Chem. Bortezomib (B-1408) from LC Laboratories. Janelia Fluor^®^ 646 HaloTag^®^ Ligand (GA1120) from Promega.

### Plasmids

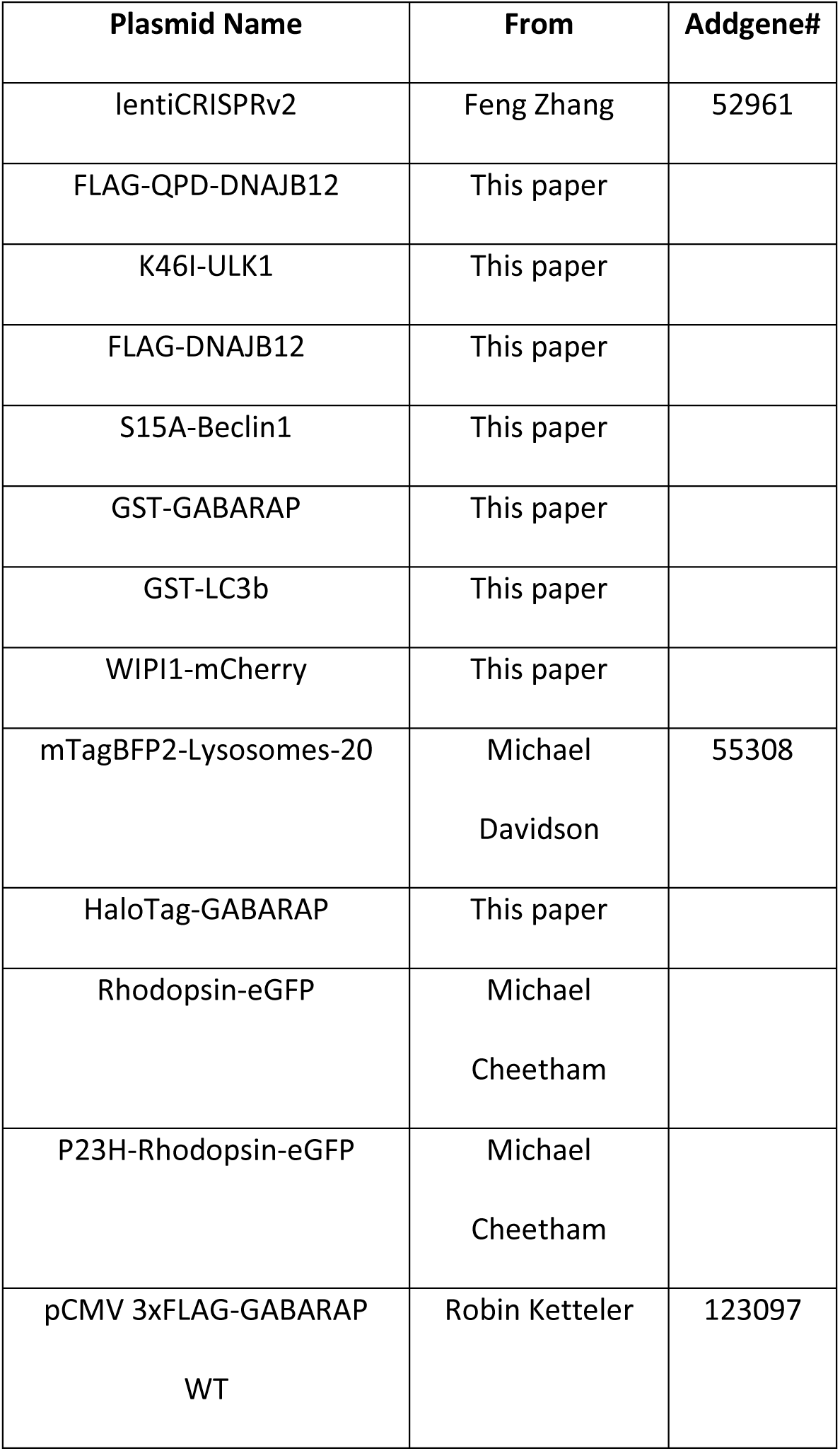
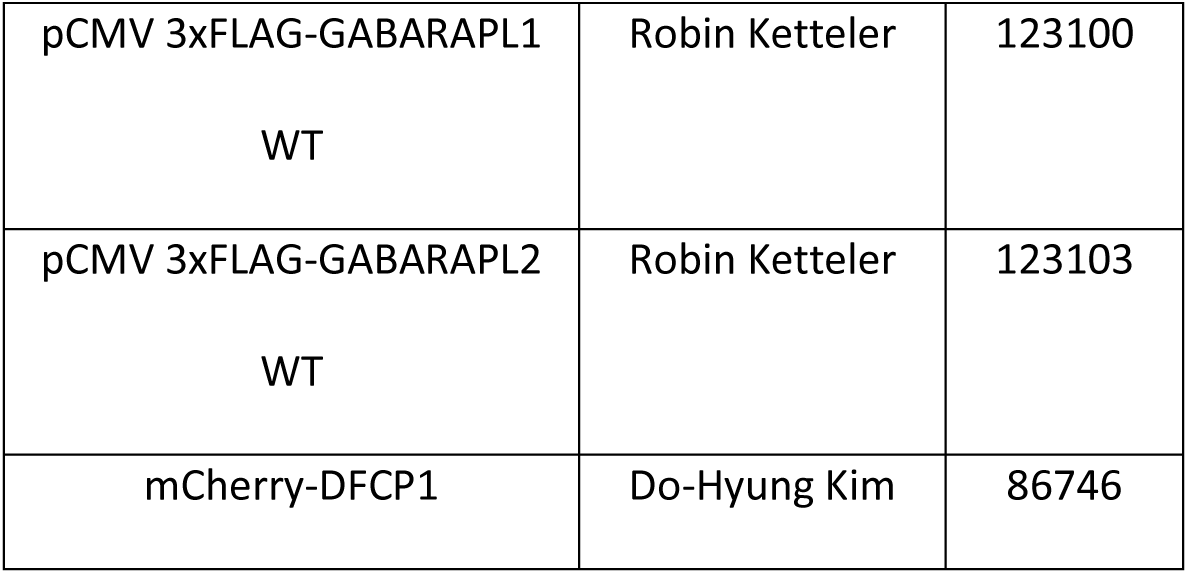

### Antibodies

| Antibody Target/Name | Host | Catalog Number | Manufacturer |
| --- | --- | --- | --- |
| anti-Mouse IgG (H+L)<br>Secondary Antibody, HRP | Goat | 31430 | ThermoFisher Scientific |
| anti-Rabbit IgG F(ab')<sub>2</sub><br>Secondary Antibody, HRP | Goat | 31461 | ThermoFisher Scientific |
| anti-Mouse IgG (H+L)<br>Highly Cross-Adsorbed<br>Secondary Antibody, Alexa<br>Fluor 488 | Goat | A11029 | ThermoFisher Scientific |
| anti-Mouse IgG (H+L)<br>Cross-Adsorbed Secondary<br>Antibody, Texas Red-X | Goat | T-862 | ThermoFisher Scientific |
| anti-Rabbit IgG (H+L) Cross-Adsorbed Secondary<br>Antibody, Texas Red | Goat | T-2767 | ThermoFisher Scientific |
| anti-Mouse IgG (H+L)<br>Highly Cross-Adsorbed<br>Secondary Antibody, Alexa<br>Fluor Plus 647 | Goat | A32728 | ThermoFisher Scientific |
| anti-Rabbit IgG (H+L) Highly<br>Cross-Adsorbed Secondary<br>Antibody, Alexa Fluor Plus<br>647 | Goat | A32733 | ThermoFisher Scientific |
| anti-Mouse IgG (H+L)<br>Highly Cross-Adsorbed<br>Secondary Antibody, Alexa<br>Fluor Plus 594 | Goat | A32742 | ThermoFisher Scientific |
| anti-Rabbit IgG (H+L) Highly<br>Cross-Adsorbed Secondary<br>Antibody, Alexa Fluor Plus<br>594 | Goat | A32740 | ThermoFisher Scientific |
| anti-Rabbit IgG (H+L) Cross-Adsorbed Secondary<br>Antibody, Alexa Fluor 405 | Goat | A31556 | ThermoFisher Scientific |
| FAM134b | Rabbit | NBP2-55248 | Novus Biologicals |
| Sec62 | Rabbit | NBP1-84045 | Novus Biologicals |
| Calnexin | Rabbit | C4731 | Sigma-Aldrich |
| GABARAP/L1/L2 | Rabbit | ab109364 | Abcam |
| LC3b | Rabbit | L7543 | Sigma |
| DNAJB12 | Rabbit | 16780-1-Ap | Proteintech |
| DNAJB14 | Rabbit | 16501-1-AP | Proteintech |
| TEX264 | Rabbit | 25858-1-AP | Proteintech |
| ULK1 | Rabbit | 4773 | Cell Signaling Technology |
| CCPG1 | Rabbit | 13861-1-AP | Proteintech |
| FIP200 | Rabbit | 10043-2-AP | ProteinTech |
| eIF2a | Rabbit | 3597 | Cell Signaling Technology |
| P-eIF2a | Rabbit | 9722 | Cell Signaling Technology |
| GFP | Mouse | 11814460001 | Roche |
| Beclin | Rabbit | 3738 | Cell Signaling Technology |
| Calnexin | Rabbit | C4731 | Sigma |
| ATL3 | Mouse | Custom | Craig Blackstone |
| WIPI | Rabbit | PA5-34452 | ThermoFisher |
| Tubulin | Mouse | T4026 | Sigma |
| GAPDH | Mouse | 60004-1-Ig | ProteinTech |
| Bip | Rabbit | 3183 | Cell Signaling Technology |
| HSP70 | Rabbit | SPA-757 | Assay Designs |
| SQSTM1 (P62) | Mouse | H00008878-M01 | Abnova |
| LAMP-2($\alpha$) | Mouse | 555803 | BD Pharmingen |
| LAMP1 | Mouse | H4A3 | Developmental Studies<br>Hybridoma Bank |
| VPS34 | Rabbit | 4263S | Cell Signaling Technology |
| BiP | Rabbit | 3183 | Cell Signaling Technology |
| GFP (EM) | Rabbit | ad6556 | abcam |
| FLAG | mouse | F1804 | Sigma |
| GFP (westerns) | Mouse | 11814460001 | Roche |
| HSP70 | Rabbit | SPA-757 | Assay Designs |
| GST Antibody | Mouse | A00865 | GenScript |

### CRISPR/Cas9 Knockout Cell Line

DNAJB12-KO, FAM134b-KO, or ‘CRISPR-control’ (vector with no insert) HeLa cell lines were generated by lentiviral transduction using the lentiCRISPRv2 plasmid (a gift from Feng Zhang - Addgene plasmid #52961). Cells were selected with 3µg/mL puromycin until complete uninfected cell death occurred, then maintained at 0.5µg/mL during passaging (puromycin removed from media before used in assays). The HeLa ATG8 knockout cell lines were provided by Wade Harper, Harvard University (Vaites et al., 2018).

### CRISPR/Cas9 Sequences

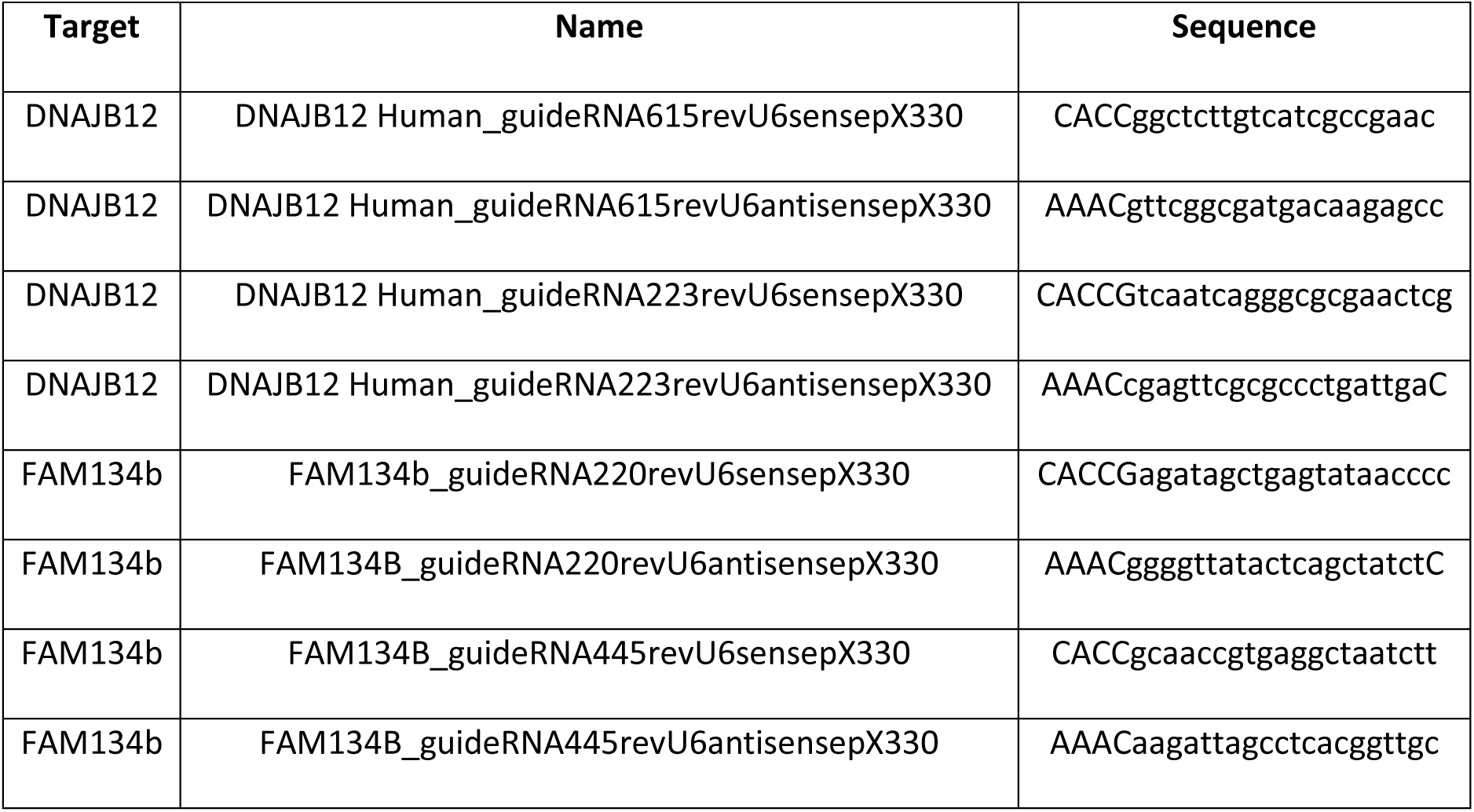

### siRNA Sequences

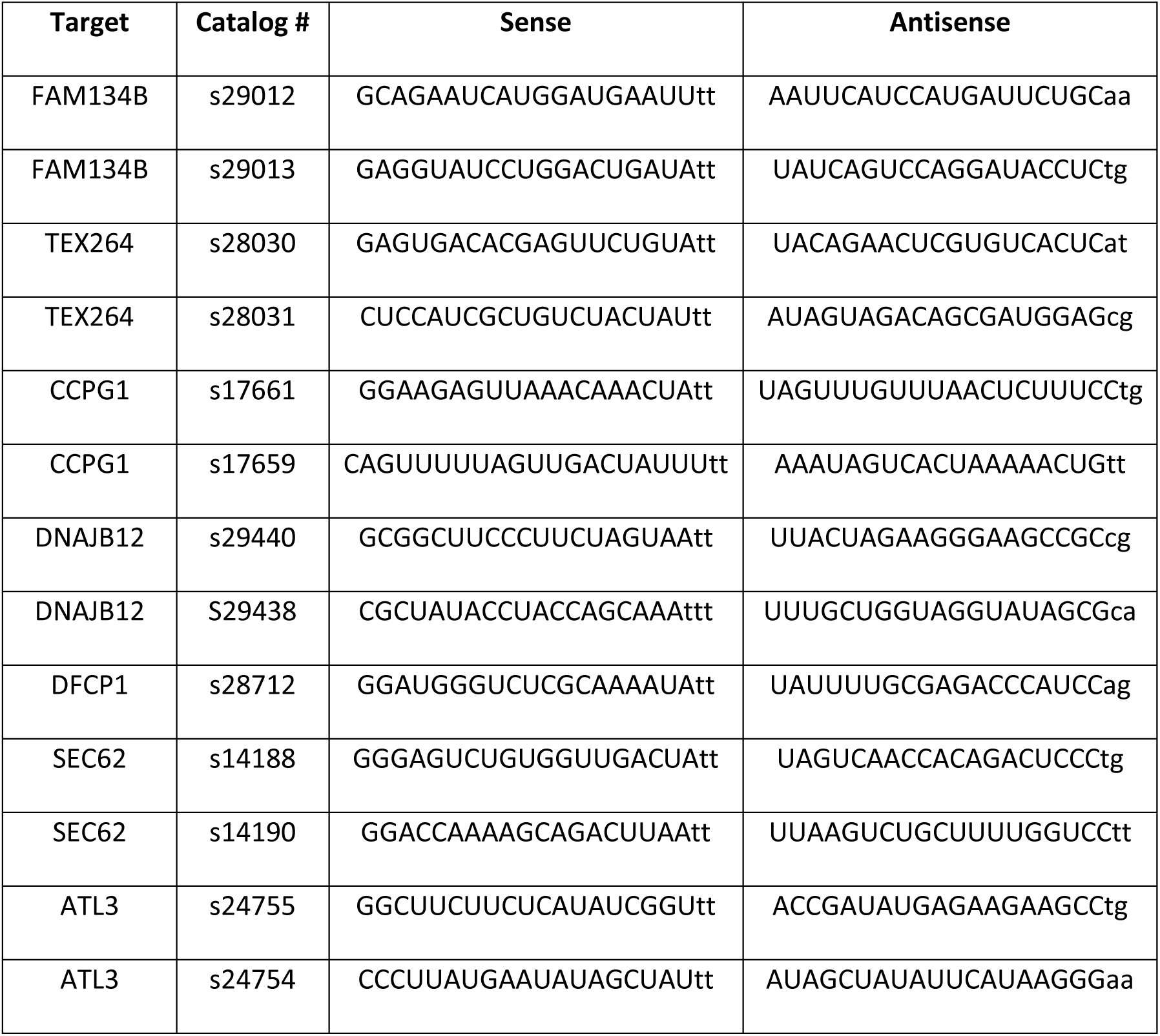
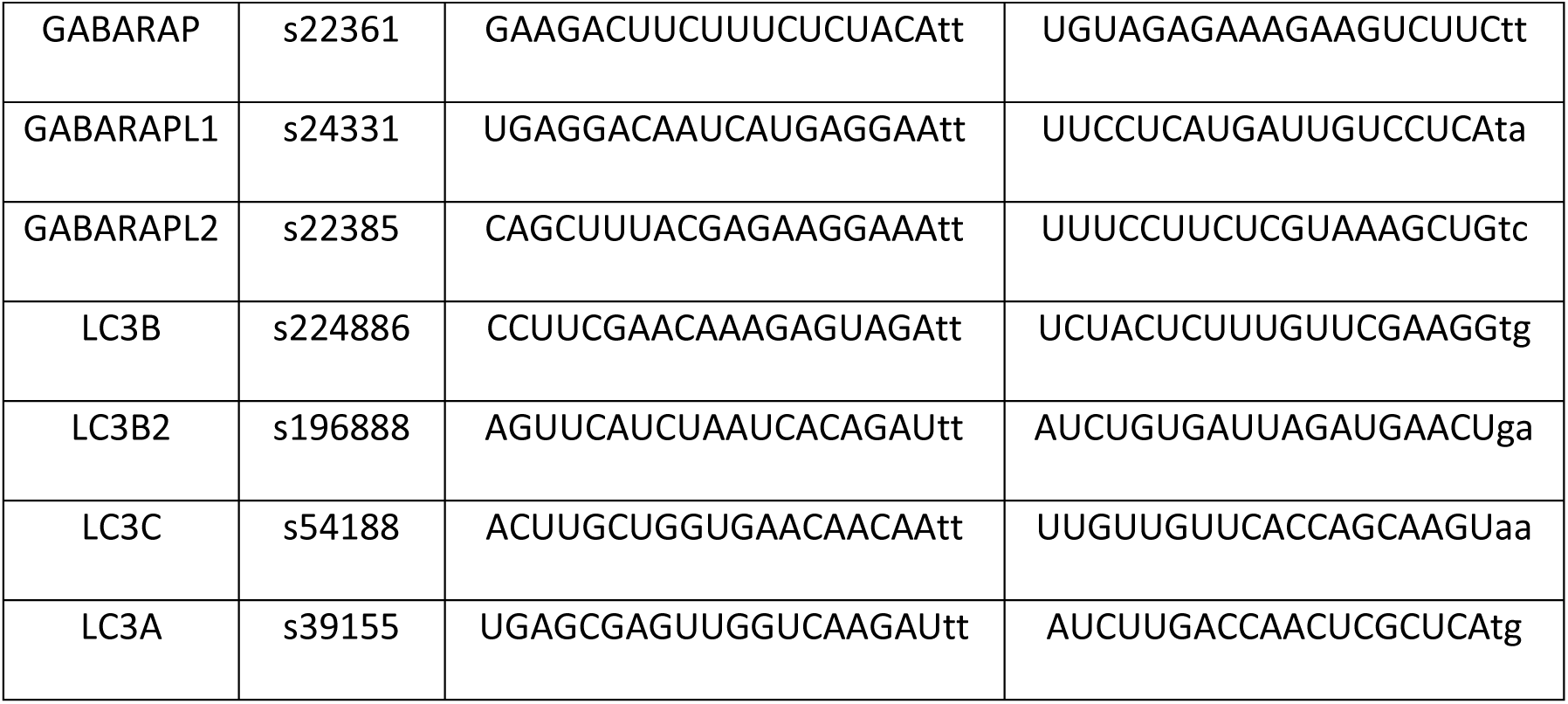

### Cell culture and transfection

African green monkey kidney cells (COS-7 ATCC CRL-1651), human embryonic kidney cells (HEK293, ATCC CRL-1573), and human cervix epithelial cells (HeLa cells ATCC-CCL2) were cultured in Dulbecco Eagle minimum enriched medium (DMEM) supplemented with 10% fetal bovine serum (Sigma-Aldrich, St. Louis, MO) and 10U/ml penicillin, 10μg/ml streptomycin (GIBCO, Carlsbad, CA) at 37°C, 5% CO_2_.

Plasmid transfections in cell lines were performed using Effectene Transfection Reagent following the product instruction. For knockdown experiments, all siRNAs were purchased from Thermo Fischer Scientific. For some target genes, two siRNAs to the same target gene were used to increase knockdown efficiency. COS-7 or HEK293 cells that were seeded into a 6-well plate at 3×10^5^ cells/well for 20 hr. were transfected at five pmol per well of siRNA with Lipofectamine RNAiMAX. Twenty hours post-transfection, the cells were split again to the appropriate density for transfection of plasmids using Effectene Transfection Reagent. The following drug treatments were used at these concentrations for 6 hours unless otherwise indicated: 100nM Bafilomycin A1; 10µM Bortezomib.

### Protein sample preparation for SDS-PAGE and Western blotting

COS-7 cells or HEK293 cells were harvested from plates with citric saline (135mM KCl, 15 mM Sodium Citrate). Cell pellets were washed in PBS, then resuspended in PBST lysis solution (1% (v/v) Triton X-100 in phosphate-buffered saline) supplemented with 1 mM PMSF, 1x complete protease inhibitor cocktail (Roche, Basel, Switzerland). 2x Laemmli sample buffer without reducing agent was added, and the total lysates were homogenized by sonication. Protein concentrations were measured using detergent compatible DC Protein Assay Reagent, and 50 mM DTT was added before samples were resolved by SDS-PAGE and proceeded to Western blot analysis. Protein signal was detected by Clarity™ western ECL substrate and visualized by LAS680 imager (GE Life Sciences, Pittsburgh, USA). Images were analyzed by ImageJ (NIH, USA) and quantified with ImageQuant (GE Life Sciences, Pittsburgh, USA).

### Co-immunoprecipitation assay

HEK293 or COS-7 cell lysates in PBST solution (1% Triton X-100 in PBS) were centrifuged at 100,000 g, at 4 °C, for 10 min to remove any insoluble fraction. Protein concentrations were measured using DC Protein Assay, and samples of an equal amount of proteins were used for immunoprecipitation analysis. To conduct pull-downs, indicated antibodies cross-linked to Protein G agarose beads (Invitrogen) were incubated with cleared lysate for 1 hr. at 4 °C. Beads were washed 3x with PBST lysis buffer. Bound proteins were eluted from beads via incubation with 2x SDS-PAGE sample buffer without reducing agent at 37°C for 15 min. Anti-FLAG M2 beads (Sigma-Aldrich, St. Louis, USA) were used to pull-down proteins interacting with FLAG-tagged proteins. After washing, proteins bound were eluted with FLAG peptide. Eluted proteins were subjected to SDS-PAGE and Western blot analysis as described above.

### Triton X-100 solubility assay

Harvested cell pellets were lysed in PBST (1% Triton X-100 in PBS) and separated into soluble and insoluble fractions by centrifugation at 20,000xg for 30 min at 4°C. The insoluble fraction was resuspended in a 2x sample buffer with an equal volume of soluble fraction and sonicated. Total lysates, soluble fractions, and insoluble fractions were resolved with SDS-PAGE and subjected to Western blot analysis.

### Immunofluorescence microscopy

COS-7 or HeLa cells were grown on 0.35mm* coverslips (Thermo Fisher Sci. 12541A***) in 6 well plates. After transfection and treatment, conditions with HaloTag fusion protein expression were also incubated with 100nM of Janelia Fluor^®^ 646 HaloTag^®^ Ligand (Promega GA11220) for 30min prior to fixation. Cells were fixed at −20 °C for 5 min with pre-cooled methanol. Cells were rehydrated by washing 3x in PBS and blocked with 3% BSA in PBS for 30 min, before immunostaining with various antibodies in 3% BSA in PBS for 1 hour. Goat anti-mouse or goat anti-rabbit secondary antibodies with different fluorophores were used to detect proteins of interest in 3% BSA in PBS for 20 minutes. Coverslips were washed 8x with PBS then mounted onto slides using ProLong Diamond antifade mountant without (P36970) or with DAPI (P36971) (Invitrogen), then sealed with nail polish. Slides were allowed to dry in the dark at room temperature for at least 12 hours before imaging with an Olympus ix-81 motorized inverted microscope with CellSens acquisition software (Olympus, Center Valley, PA) or LSM 880 with Airyscan with ZEN acquisition software (Zeiss, Jena, Germany).

### Live cell imaging

COS-7 cells were cultured in 35mm glass-bottom dishes (MatTek P35G-1.5-10-C). After overnight transfection with fluorescent-tagged proteins, cells were rinsed with pre-warmed FluorBrite DMEM (Gibco Life Sci.) and kept in the same medium during drug treatment and imaging. Live cell imaging was carried out using LSM 880 with Airyscan and live-cell incubation chamber (Zeiss, Jena, Germany) where indicated. For all other: Images were acquired by an inverted Olympus FV3000RS confocal laser point-scanning microscope (Olympus Corporation, Tokyo, Japan) equipped with four GaAsP detectors, resonant and galvanometer scanners attached to a fully motorized Olympus IX83 inverted stand equipped with an Olympus IX3-SSU motorized scanning stage encoded with an ultrasonic motor in the UNC Neuroscience Microscopy Core (RRID:SCR_019060), equipped with a Tokai Hit stage-top incubation chamber (model: STXG-WSKMX-SET) at 37°C with 5% CO2 and humidity using a 60x/1.4NA PLAPON oil immersion (Olympus) objective lens. A 405 nm 50 mW diode laser was used for excitation of Alexa Fluor 405, a 488 nm 20 mW diode laser was used for excitation of mEGFP, a 561 nm diode 20 mW laser was used for excitation of Alexa Fluor 568 and a 640 nm 40 mW diode laser was used for excitation of Alexa Fluor 647. A 405/488/561/640 dichroic mirror was used to acquire all channels. Laser power was adjusted between cells to maintain similar fluorescence intensity histogram between cells, with final transmittance of laser power set in the software ranging from 1.0-8.5%, with a 10% ND filter in the light path. All images were acquired in resonance scanning mode, at 16 bit depth, with channels acquired sequentially by line on GaAsP detectors, using 10x frame averaging with a pinhole size of 217 µm, a zoom of 4.62x, a frame size of 512 x 512 pixels with pixel sizes of 0.090 µm, and z-step of 0.42 µm between slices. Light emitted from the Alexa Fluor 405 channel was collected in the detector unit HSD1 (GaAsP detector) with a variable bandpass filter range of 409-484 nm and voltage (HV) set to 500 volts. Light emitted from the mEGFP channel was collected in the detector unit HSD3 (GaAsP detector) with a variable bandpass filter range of 492-557 nm and voltage (HV) set to 500 volts. Light emitted from the Alexa Fluor 568 channel was collected in the detector unit HSD2 (GaAsP detector) with a variable bandpass filter range of 575-638 nm and voltage (HV) set to 650 volts. Light emitted from the Alexa Fluor 647 channel was collected in the detector unit HSD4 (GaAsP detector) with a variable bandpass filter range of 646-746 nm and voltage (HV) set to 480 volts. Images were acquired and deconvoluted using CellSens software (Olympus) and processed with Bioimage ICY (de Chaumont, 2012) and Adobe Photoshop.

### Image Quantitation and Processing

All views of microscopy images with scale bars were made using Bioimage ICY (v2.2.1.0) (de Chaumont, 2012) with the ‘Scale Bar’ (Thomas Provoost - http://icy.bioimageanalysis.org/plugin/scale-bar/) (v3.2.0.0) plugin. Signal profiles of images were generated using a line ROI drawn in Bioimage ICY then ‘Plot Profile’ was run using the built-in ImageJ within ICY. Profiles were normalized to the maximum intensity. For live-cell plot profiles, the lowest intensity was subtracted within each channel, then normalized to the maximum intensity.

Statistical analysis of signal co-localization was achieved through the calculation of Pearson’s correlation coefficients (R). For images with an R value inscribed in the image, an ROI was manually drawn and cropped around the cell of interest or inset, followed by extraction of the in-focus plane using Bioimage ICY. The ‘Coloc 2’ plugin for FIJI/ImageJ was then used to calculate Pearson’s correlation coefficients between eGFP-Rhodopsin and different sets of markers (Schindelin, 2012). For images where Pearson’s R was calculated within identified objects, a custom CellProfiler pipeline was developed to identify LAMP1/2 objects within cells expressing Rhodopsin, then calculate Pearson’s R within each object for any given marker. Statistical analysis a plot generation was performed in Prism GraphPad 9 (San Diego, CA). The fractional intensity analysis utilized the ‘MeasureObjectIntensityDistribution’ function in CellProfiler (McQuin, 2018).

### GST-GABARAP and GST-LC3 purification

GST_GABARAP was expressed in *E. coli* strain BL21(DE3) and purified as follows: pellet from 300ml culture were harvested after a 3 hours induction with 0.2mM IPTG and lysed by sonication in buffer PBS (140mM NaCl,2.7mMKCl, 10mM Na2HPO4, 1.8mM KH2PO4 pH 7.4), 2mM DTT, 0.1mM PMSF. The high-speed supernatant was loaded onto GSTrap 4B column equilibrated with PBS. Bound GST-GABARAP was eluted by 50mM Tris-HCl 20mM reduced glutathione pH8.0. After final eluted dialyzed against 20mM HEPES pH 7.4, 150mM NaCl, 2mM DTT, 0.1mM PMSF. Aliquots of protein were frozen in liquid nitrogen and store at −80C until used.

### GST-GABARAP and GST-LC3 pulldown assays

HEK293 cells were lysed at 4°C in PBS buffer 1% Triton X-100, Protease Inhibitor Mixture Tablets Complete (Roche Applied Science), and 1 mM PMSF. High speed supernatant was incubated with GST-GARARAP or GST-LC3 30minutes at 4°C, then added Glutathione Sepharose (GE 17-5132-01) for 30 minutes at 4°C. After washing with lysis buffer, the samples eluted with 2× SDS sample buffer (100 mM Tris-HCl, pH 6.8, 4% SDS, 0.05% bromophenol blue, and 20% glycerol)

### Electron Microscopy

Cells were harvested 24 hours post transfection with 0.05% Trypsin-EDTA (Gibco) from a 6-well plate and reseeded on Nunc Permanox chamberslides. Cells were allowed to grow for another 24 hours and then were fixed with 2% paraformaldehyde/0.5% glutaraldehyde in 0.15 M sodium phosphate buffer, pH 7.4, for 1 hour. Following free aldehyde inactivation with 0.2 M glycine in 0.15 M sodium phosphate buffer (PB), cells were permeabilized with 0.1% saponin in PB for one hour, and incubated in a 1:100 dilution of Rabbit anti-GFP (Cell Signaling, #2956) overnight at 4°C. After buffer washes, samples were incubated in secondary antibody, 1:100 goat anti-rabbit IgG 0.8nm immunogold (Aurion, Electron Microscopy Sciences), for 16 hours at 4°C. The cells were post-fixed in 2% glutaraldehyde in PB, and silver-enhanced for 90 minutes using an Aurion R-Gent SE-EM Silver Enhancement Kit. The monolayers were post-fixed in 0.1% osmium tetroxide, dehydrated in ethanol and embedded in Polybed 812 epoxy resin (Polysciences, Inc., Warrington, PA). 80nm ultrathin sections were cut, mounted on copper grids, and post-stained with 4% uranyl acetate and lead citrate. Sections were observed using a JEOL JEM 1230 transmission electron microscope (JEOL USA, Inc. Peabody, MA) operating at 80kV and images were taken using a Gatan Orius SC1000 CCD camera with Digital Micrograph 3.11.0 (Gatan, Inc., Pleasanton, CA).

## Supplementary Materials

**Figure 2 Supplement.**
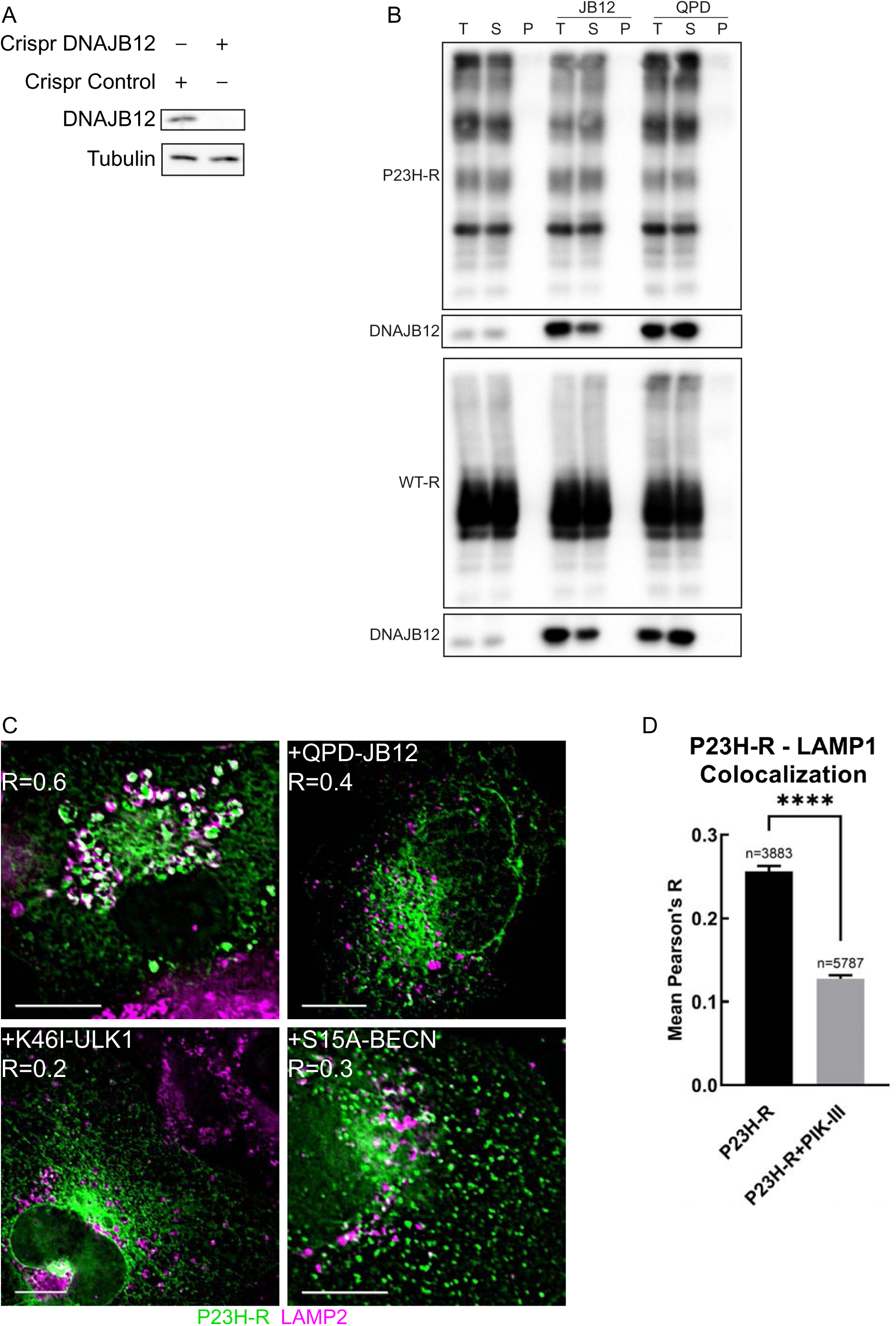
(A) Western blot for DNAJB12 in CRISPR-Control/DNAJB12 cell lines utilized in Figure 2B (B) Total/pellet/supernatant assay to determine aggregation propensity: COS-7 cells expressing P23H-R and DNAJB12/QPD lysed and rotated with 1% Triton-X for 1 hour, followed by centrifugation and harvest of supernatant and pellet fractions; Western blot of each fraction or total (pre-centrifugation) (C) Fixed-cell epifluorescent microscopy of COS-7 cells transiently transfected with P23H-R and dominant-negative forms of DNAJB12 (QPD-JB12), ULK1 (K46I-ULK1), or Beclin1 (S15A-BECN) (D) Bar plot showing mean of Pearson’s R between P23H-R and LAMP1 within automatically identified vesicles (COS-7 cells transiently transfected with P23H-R then treated ± 2.5µM PIK-III (inhibitor of VPS34) followed by fixation and immunostaining for LAMP1) Scale bars: 10µm

**Figure 3 Supplement.**
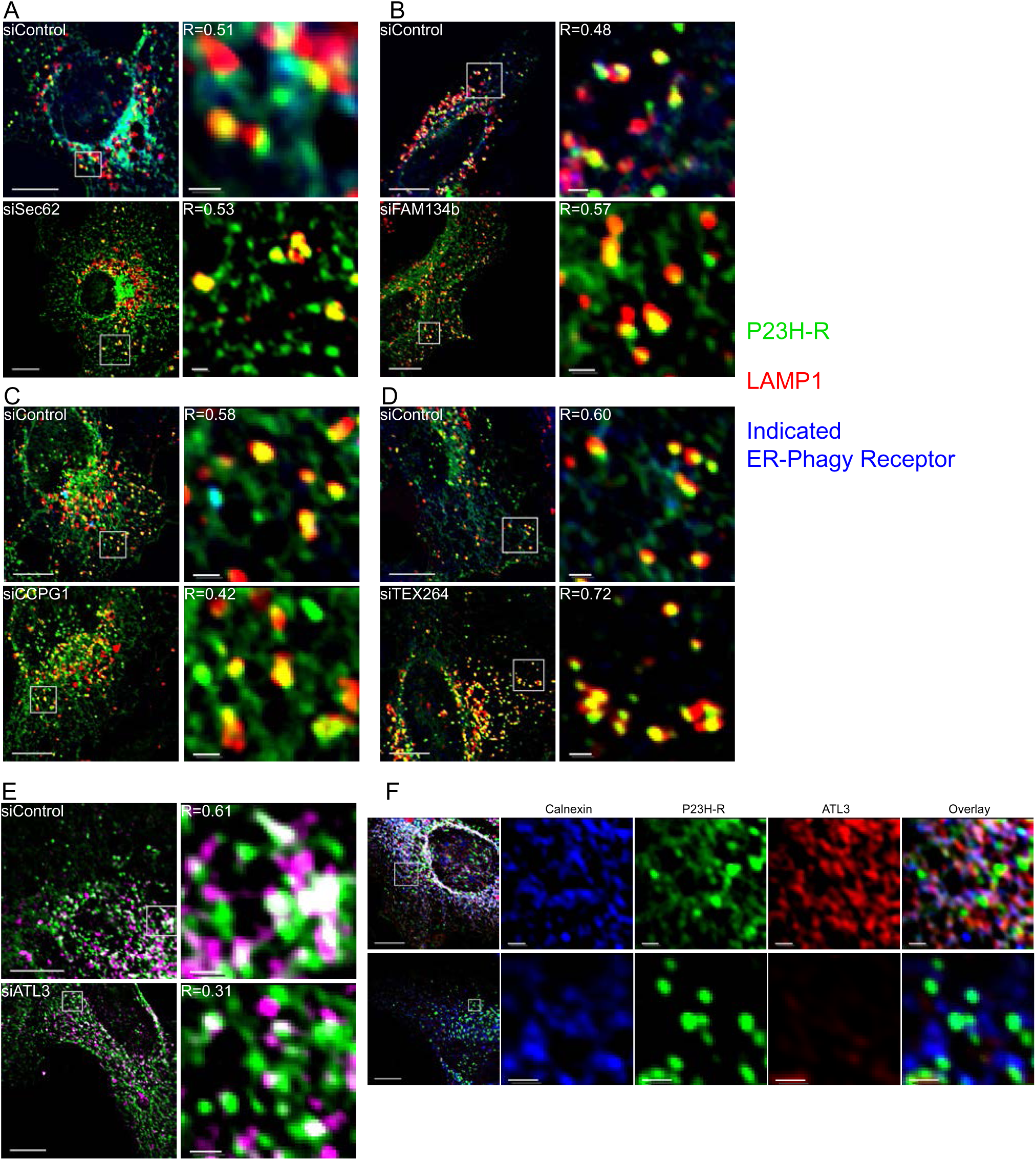
(A-E) Fixed-cell epifluorescent microscopy of COS-7 cells transiently transfected with P23H-R after siRNA knockdown of indicated ER-phagy receptor. R indicates Pearson’s R for Rhodopsin-LAMP1 colocalization (F) Additional staining of siATL3 + P23H-R with calnexin immunofluorescence Scale bars: whole cell views = 10µm; insets = 1µm

**Figure 4 Supplement.**
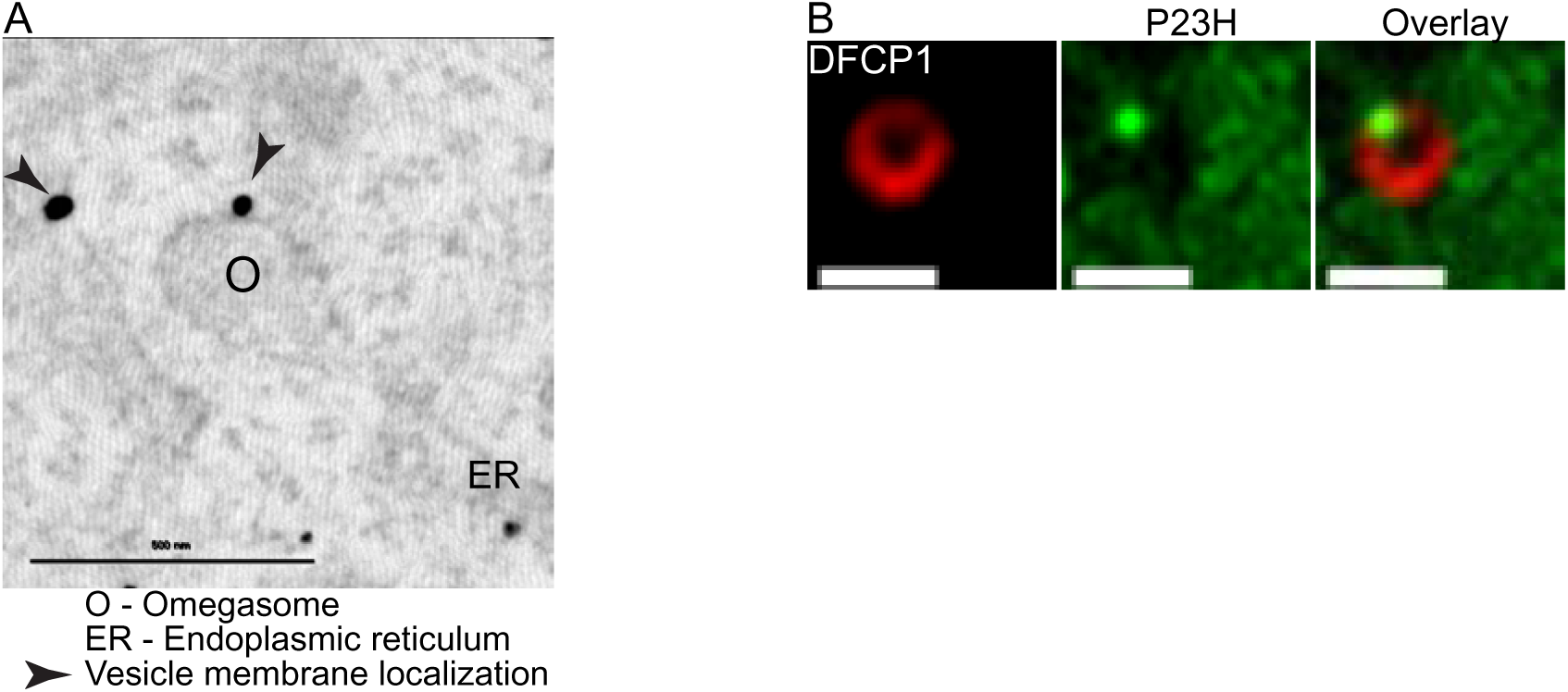
(A) Transmission EM image of anti-GFP immunogold staining in COS-7 cells transiently transfected with P23H-R. Scale bar = 500nm (B) Live-cell confocal microscopy of COS-7 cells expressing P23H-R and either WIPI1-mCherry or DFCP1-mCherry. Scale bar = 1µm

**Figure 5 Supplement.**
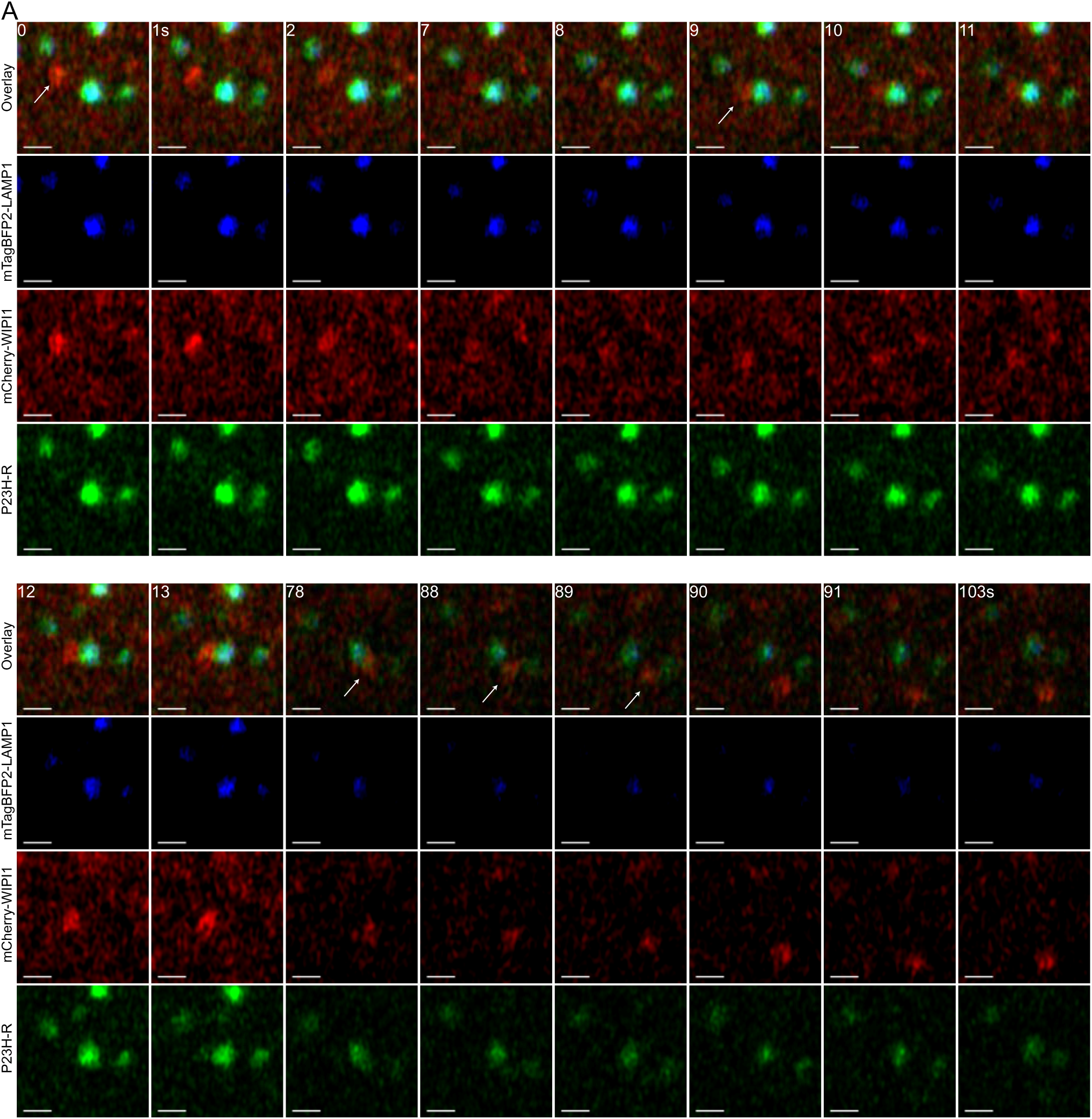
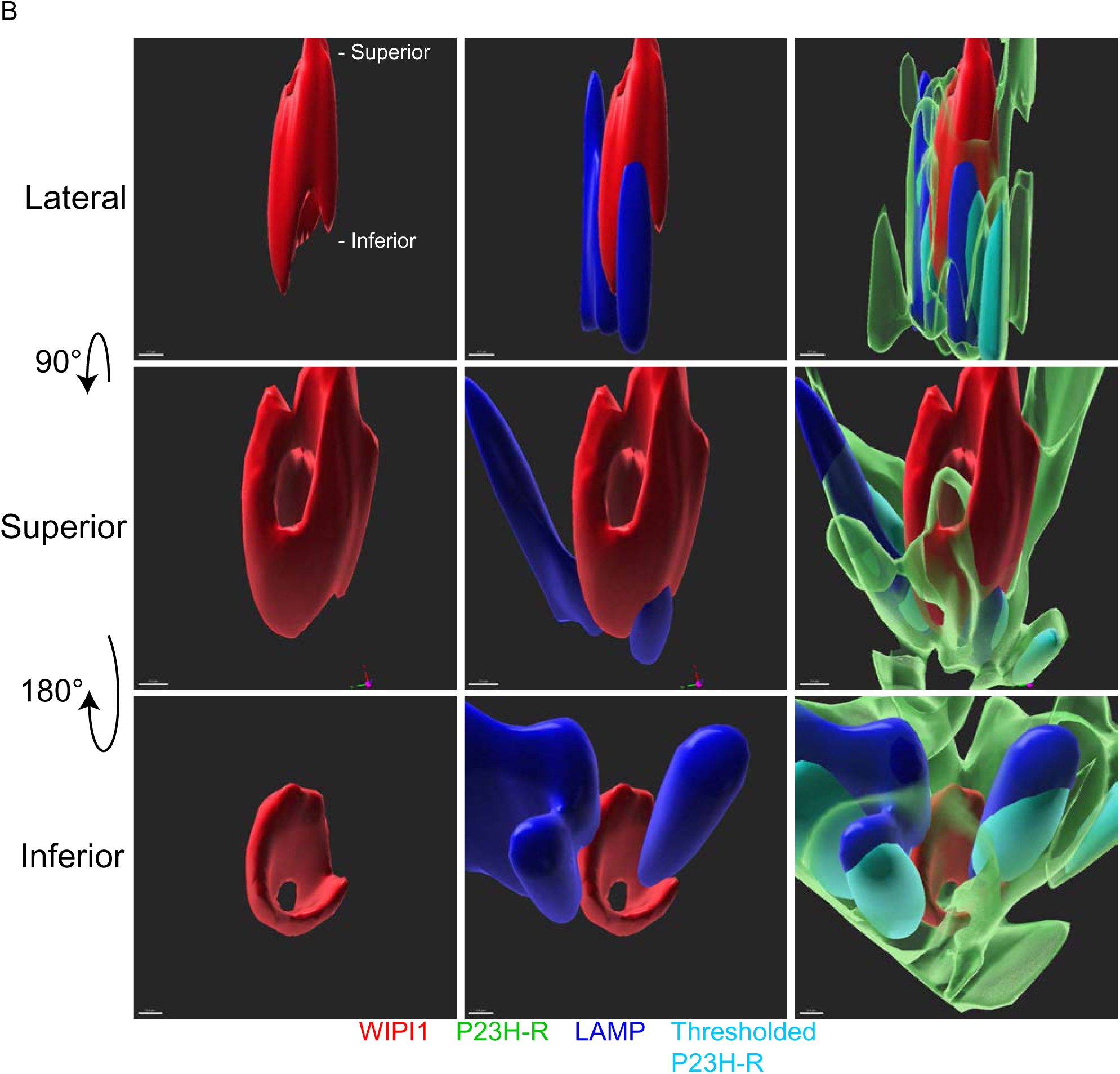
(A) Airyscan time-lapse of COS-7 cells expressing P23H-R, mTagBFP2-LAMP1, and WIPI1-mCherry. Scale bar = 1µm (B) Alternate views of *5C*. Lateral scale bar = 0.7µm; superior and inferior scale bars = 0.5µm

**Figure 6 Supplement.**
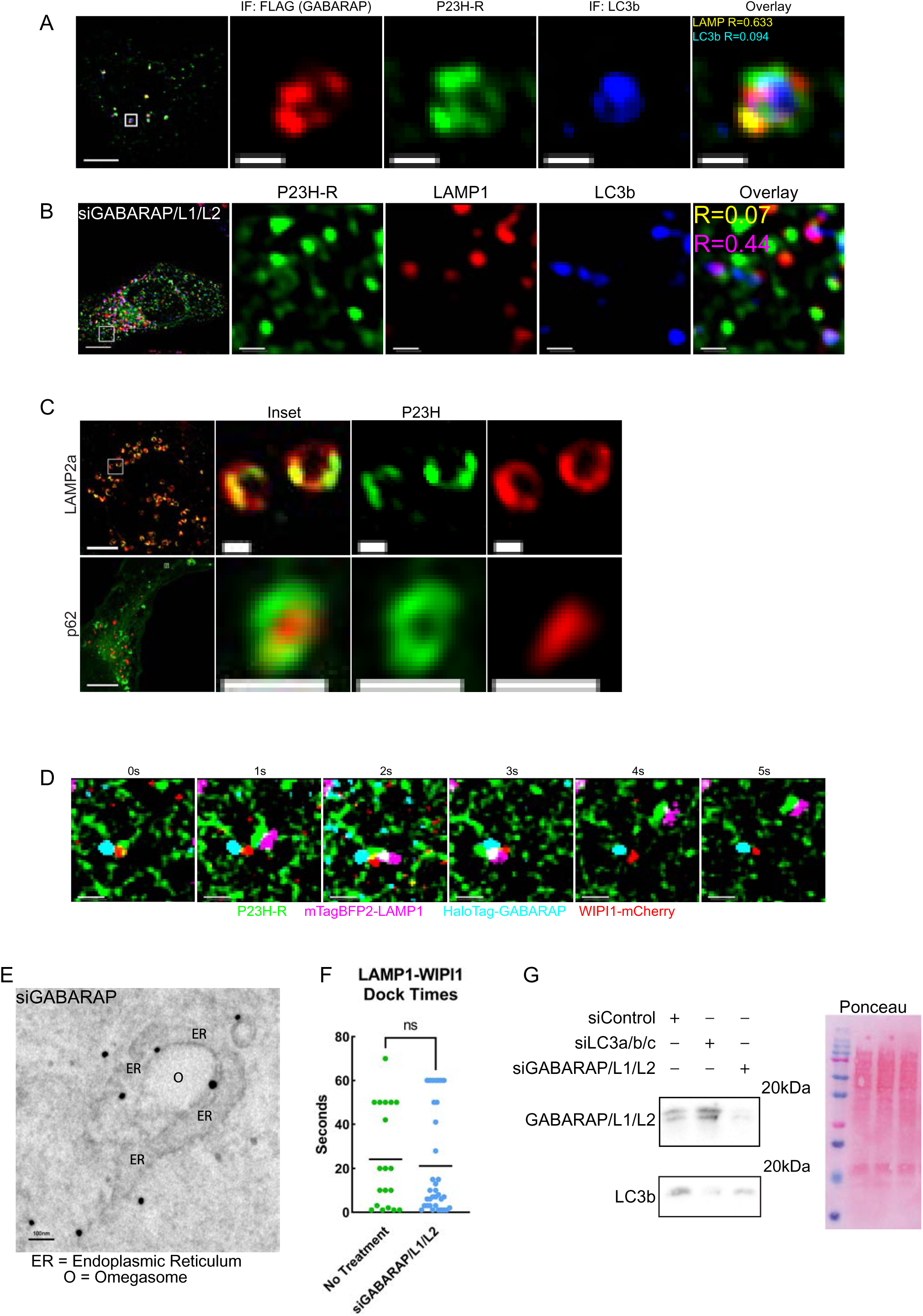
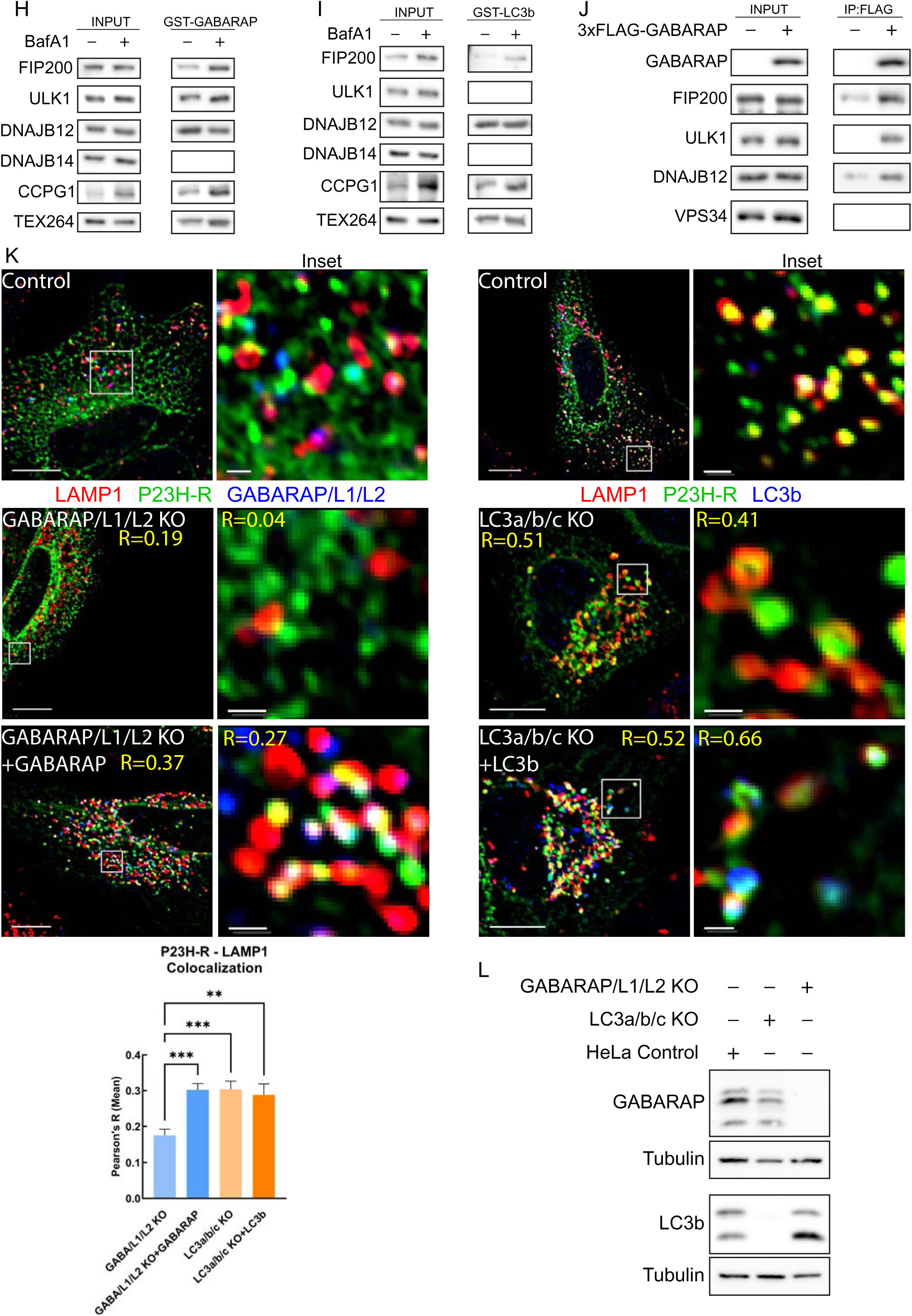
(A) Fixed-cell epifluorescent microscopy of COS-7 cells expressing P23H-R and 3xFLAG-GABARAP, treated with 100nM BafA1, and immunostained for FLAG and LC3b. For inset overlay, top value (yellow) refers to GABARAP and P23H-R Pearson’s R; bottom value (cyan) refers to LC3b and P23H-R Pearson’s R (B) Fixed-cell epifluorescent microscopy of COS-7 cells expressing P23H-R after siGABARAP/L1/L2 from *6A* but with immunofluorescence staining for LC3b and LAMP1. In inset overlay: top value (yellow) refers to LAMP1 and P23H-R Pearson’s R; bottom value (magenta) refers to LC3b and LAMP1 Pearson’s R (C) Fixed-cell Airyscan images of COS-7 cells expressing P23H-R, treated with 100nM BafA1, and stained for either LAMP2a or p62 (D) View of *6D* without manual channel-offset correction (E) TEM image of anti-GFP immunogold staining in siGABARAP/L1/L2 COS-7 cells transiently transfected with P23H-R. Scale bar = 100nm (F) Dot plot of observed LAMP1-WIPI1 dock times with unpaired t-test - line indicates mean. ‘No Treatment’ vs ‘siGABARAP/L1/L2’: n=19 and 34, Range=69 and 59, Mean=24.26 and 21.15, SEM=5.269 and 2.076 respectively (G) Western blot of COS-7 cells for endogenous GABARAP/L1/L2 or LC3b after siLC3a/b/c or siGABARAP/L1/L2 (H) Western blot of GST pull-down in HEK-293 cells transiently transfected with GST-GABARAP ± 100nM Bafilomycin A1 (BafA1) (I) Western blot of GST pull-down in HEK-293 cells transiently transfected with GST-LC3b ± 100nM Bafilomycin A1 (BafA1) (J) Western blot of FLAG pull-down in HEK-293 cells with and without transient transfection of 3xFLAG-GABARAP (K) Top: Fixed-cell epifluorescent microscopy of HeLa cells with or without GABARAP/L1/L2 or LC3a/b/c knockout cell lines (KO conditions treated with BafA1) ±GABARAP or LC3b respectively. Bottom: Bar graph of mean Pearson’s R values between Rhodopsin and LAMP1 with one-way ANOVA and Dunnett’s multiple comparisons test (n=321 cells). Error bars for SEM (L) Western blot of cells used in *K* ‘GABARAP/L1/L2’ includes GABARAP, siGABARAP-L1, siGABARAP-L2 concurrently; ‘LC3abc’ includes LC3a, LC3b, and LC3c concurrently **P < 0.01, ***P < 0.001 Scale bars: whole cell views = 10µm; insets = 1µm

